# Cryo-EM structures of the MnmE-MnmG complex reveal large conformational changes and provide new insights into the mechanism of tRNA modification

**DOI:** 10.1101/2024.11.29.625835

**Authors:** Laila Maes, Ella Martin, David Bickel, Siemen Claeys, Wim Vranken, Marcus Fislage, Christian Galicia, Wim Versées

**Affiliations:** Structural Biology Brussels, Vrije Universiteit Brussel, Pleinlaan 2, 1050 Brussels, Belgium; VIB-VUB Center for Structural Biology, VIB, Pleinlaan 2, 1050 Brussels, Belgium; Interuniversity Institute of Bioinformatics in Brussels, ULB-VUB, Brussels, Belgium

## Abstract

MnmE and MnmG form a conserved protein complex responsible for the addition of a 5-carboxymethylaminomethyl (cmnm^5^) group onto the wobble uridine of several tRNAs. Within this complex, both proteins collaborate intensively to catalyze a tRNA modification reaction that involves glycine as a substrate in addition to three different cofactors, with FAD and NADH binding to MnmG and methylenetetrahydrofolate (5,10-CH_2_-THF) to MnmE. Without structures of the MnmEG complex it remained enigmatic how these substrates and co-factors can be brought together in a concerted manner. Prior SAXS data suggested that the MnmE (α_2_) and MnmG (β_2_) homodimers can adopt either an α_2_β_2_ or α_4_β_2_ complex, depending on the nucleotide state of MnmE. Here, we report the cryo-EM structures of the MnmEG complex in the α_2_β_2_ and α_4_β_2_ oligomeric states. These structures reveal that MnmE undergoes large conformational changes upon interaction with MnmG, resulting in an asymmetric MnmE dimer. In particular, the functionally important C-terminal helix of MnmE relocates from the 5,10-CH_2_-THF binding pocket of MnmE to the FAD binding pocket of MnmG, thus suggesting a mechanism for the transfer of an activated methylene group from one active site to the other. Together, these findings provide crucial new insights into the MnmEG-catalyzed reaction.

**Graphical abstract:** 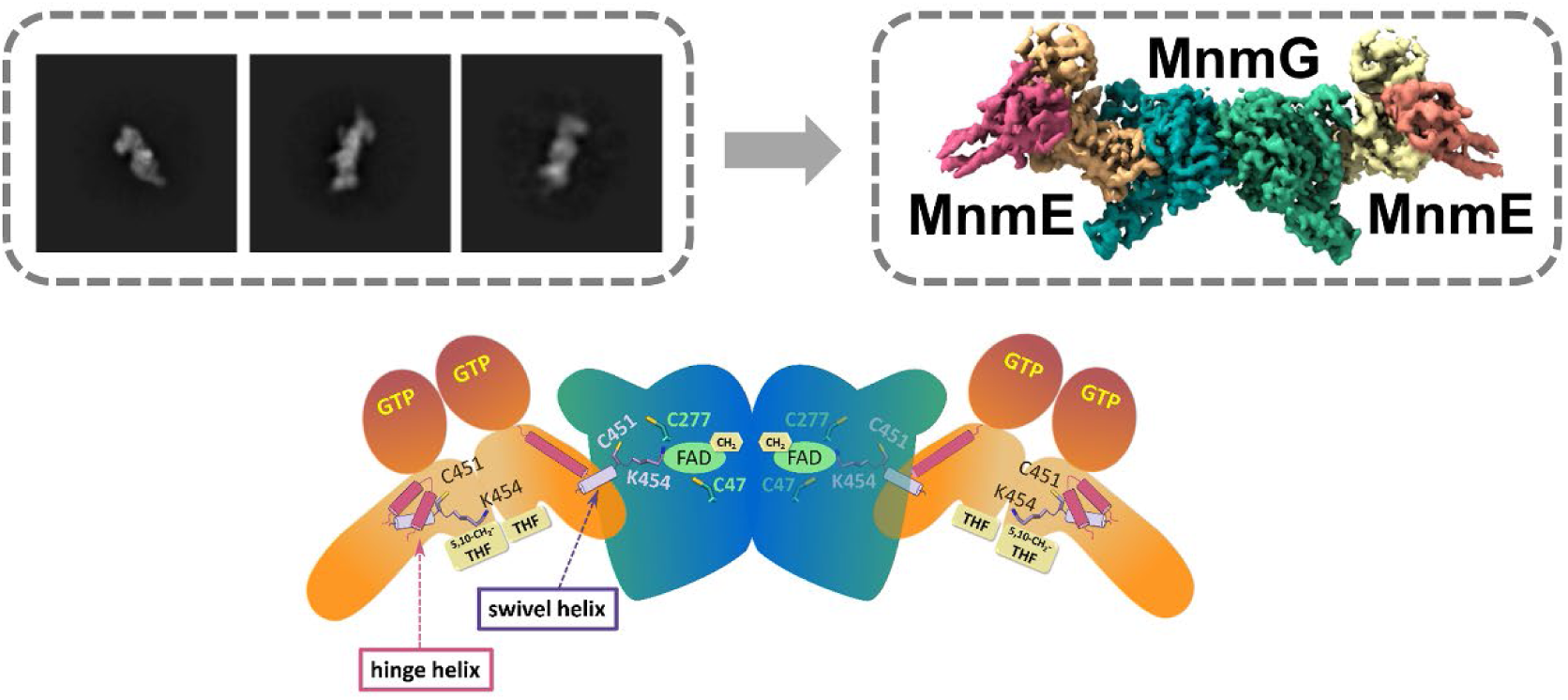

## Introduction

Transfer RNA (tRNA) is a key component in the translation process, acting as an adaptor between mRNA and the elongating peptide in the ribosome. As part of their maturation process, tRNAs are subject to a wide variety of enzyme-catalyzed post-transcriptional modifications, which are crucial for ensuring tRNA stability, aminoacylation capability and correct codon-anticodon pairing [1–4]. The physiological importance of these modifications is illustrated by the plethora of human diseases that are associated with mutations in genes coding for tRNA-modifying enzymes and the associated impairment in proper tRNA modification, including mitochondrial diseases, neurological disorders and cancer [5–8]. Additionally, the dynamic control of the tRNA modification status has been proposed as a mechanism to regulate responses to environmental stresses or to changes in metabolite levels [9–12].

While modifications occur throughout the complete tRNA molecule, a hotspot for complex modification is the wobble base on position 34 of the tRNA, and in particular the wobble uridine (U_34_) [13,14]. A universally conserved type of U_34_ modifications are the 5-methyluridine-derived (xm^5^U) modifications that ensure proper reading of split codon boxes ending with adenine or guanine [15]. In bacteria, modification of U_34_ at the C5 position with either a methylaminomethyl (mnm) or carboxymethylaminomethyl (cmnm) group is initiated by a transient enzyme complex of MnmE and MnmG [16,17]. Together these enzymes catalyze the incorporation of either an aminomethyl (nm) or carboxymethylaminomethyl (cmnm) group, using 5,10-methylenetetrahydrofolate (5,10-CH_2_-THF) and, respectively, ammonium or glycine as substrates [16,18]. The homologues of MnmE and MnmG in eukaryotes are localized to the mitochondria. Interestingly, while the corresponding yeast proteins, MSS1 and MTO1, incorporate a cmnm^5^U_34_ group in mitochondrial tRNAs, the human enzymes, GTPBP3 and MTO1, use taurine instead of glycine resulting in a τm^5^U_34_ modification [19]. In contrast, in the cytoplasm of eukaryotes this modification does not occur, and, instead, a 5-methoxycarbonylmethyl (mcm) group is being incorporated at the C5 position of U_34_ of 12 tRNAs by the elongator complex [5,20]. Mutations in the genes coding for GTPBP3 and MTO1 lead to severe mitochondrial diseases, including cardiomyopathy, encephalopathy, lactic acidosis, optic neuropathy and cognitive disability [21–23]. On the other hand, loss of either MnmE or MnmG in pathogenic bacteria, including *Pseudomonas*, *Salmonella* and *Aeromonas* species, leads to a decrease in virulence [24–26].

Crystal structures of the individual MnmE and MnmG proteins from different species have been solved many years ago [17]. MnmE is a homodimeric protein of 50-55 kDa subunits, with each protomer composed of three discernable domains: an N-terminal domain required for constitutive dimerization and binding of 5,10-CH_2_-THF, a helical domain and a Ras-like GTPase domain (G domain) that is inserted in the helical domain [27,28] (**Fig. 1A**). The α-helical domain contains a conserved C-terminal FC(V/I/L)GK motif, of which the cysteine residue (C451 in *E. coli* MnmE) is crucial for the tRNA modification reaction. Based on its biochemical properties and the observation that the isolated G domains dimerize upon binding to GDP-AlFx and potassium, MnmE has been classified as a GTPase activated by nucleotide-dependent dimerization (GAD) [29–31]. In contrast to classical Ras-like GTPases, MnmE shows a relatively low affinity for GTP/GDP, while its intrinsic GTPase activity is triggered by potassium-mediated dimerization and cross-activation of the two adjacent G domains within the MnmE homodimer. Nevertheless, so far, no structures have been solved of any full-length MnmE protein in this “closed G-domain state”. The MnmG structures also display the protein as a homodimer of approximately 70kDa subunits [32–34]. Each subunit consists of an FAD-binding domain, two insertion domains, and a helical domain (**Fig. 1A**). Within the MnmEG complex, MnmG is responsible for binding the cofactors FAD and NADH, and it is also the main driver for binding of the tRNA substrate. *E. coli* MnmG contains four cysteine residues, of which two (C47 and C277) were shown to be important for the tRNA modification reaction [34].

**Figure 1.**
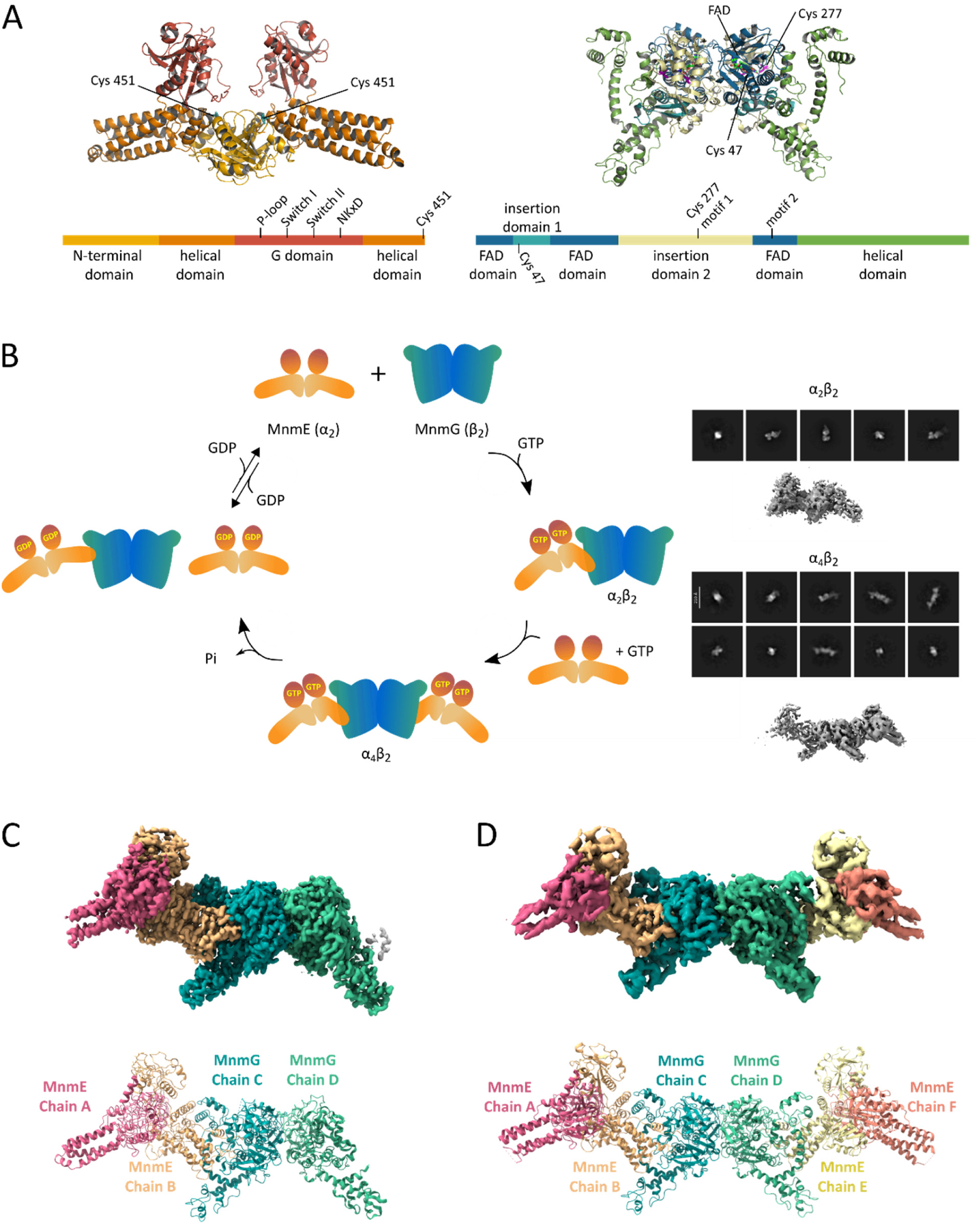
Cryo-EM structures of the MnmE-MnmG α_2_β_2_ and α_4_β_2_ complexes. (**A**) Alphafold2 models of the *E. coli* MnmE (α_2_, left) and MnmG (β_2_, right) homodimers colored by domain, and the corresponding schematic representation of their domain arrangement. (**B**) SAXS and SEC-MALS analyses previously showed that MnmE and MnmG cycle between α_2_β_2_ and α_4_β_2_ oligomeric states concomitant with MnmE GTP binding and hydrolysis [38]. While structures of MnmE and MnmG from several organisms have been solved previously [27,28,32–34], the current Cryo-EM analyses provide the first structural information on the α_2_β_2_ and α_4_β_2_ complexes. On the right representative 2D classes and initial maps of both oligomeric states are shown. (**C**) Cryo-EM map of the MnmE-MnmG α_2_β_2_ oligomeric state at 3.2 Å (top) and the corresponding structural model (bottom). The map is colored according to the different chains within the MnmEG structural model as indicated below. (**D**) Cryo-EM map of the MnmE-MnmG α4β_2_ oligomeric state at 4.1 Å (top) and the corresponding structural model (bottom). The map is colored according to the different chains within the MnmEG structural model as indicated below. The maps shown in (C) and (D) were sharpened using DeepEMhancer and EMready for visualization.

To perform the tRNA-modification reaction, MnmE and MnmG have to collaborate within an MnmEG complex, and several mechanisms for the MnmEG-catalyzed reaction have been proposed [16,18,35–37]. Two recent studies from Bommisetti and colleagues convincingly showed that FADH_2_ is used to transfer the methylene group of the one-carbon donor 5,10-CH_2_-THF onto the C5 atom of tRNA U_34_ in a covalent manner via an FADH iminium intermediate [36,37]. Subsequently, the methylene group would undergo addition of the nucleophilic substrate, glycine, to form the cmnm moiety. Despite this progress in our understanding of the mechanism, many open questions remain in the absence of structural information regarding the architecture of the MnmEG protein complex. In particular, it remains enigmatic how the different substrates and co-factors are brought together in a concerted manner during the tRNA modification reaction, with 5,10-CH_2_-THF binding to MnmE, and tRNA, FAD and NADH binding to MnmG.

The first low resolution structural insight into the overall arrangement of the protein complex formed by MnmE and MnmG was provided by small angle X-ray scattering (SAXS) more than ten years ago [38]. This study showed an intriguing change in oligomeric state of the MnmEG complex during its GTPase cycle and depending on the bound nucleotide. In the GDP-bound state, MnmE and MnmG form an asymmetric, L-shaped complex with α_2_β_2_ stoichiometry (i.e. one MnmE dimer bound to one MnmG dimer). However, upon binding of K^+^ and GTP, MnmE and MnmG adopt an elongated α_4_β_2_ complex, where one MnmG dimer is flanked on each side by an MnmE dimer (**Fig. 1B**). The GTP binding- and hydrolysis-driven interconversion between the two states happens on a time scale that is physiologically relevant, suggesting it to be an integral part of the MnmEG catalytic cycle. Nevertheless, while these SAXS models provide an overall shape of the MnmEG complex, the resolution is largely insufficient to draw any conclusions regarding the details of the interactions between MnmE and MnmG within the complex or regarding any conformational changes that might occur upon complex formation.

In the current study, we have used single particle cryo-electron microscopy (cryo-EM) to solve the structure of the MnmEG complex bound to GppNHp, showing the occurrence of both the α_2_β_2_ and α_4_β_2_ oligomeric states. These structures reveal large and unanticipated conformational changes in the subunit of MnmE that interact with MnmG. An important finding is a large relocation of the C-terminal helix of this subunit, from the 5,10-CH_2_-THF-binding pocket of MnmE toward the FAD binding pocket of MnmG. The potential functional implications with regard to the tRNA modification reaction are discussed.

## Materials and Methods

### Protein expression and purification

*E. coli* MnmE and MnmG were expressed and purified as previously described [38]. In brief, protein expression was initiated from *E. coli* BL21(DE3) cells transformed with either the pET20b vector encoding wild-type MnmE preceded by an N-terminal His-tag and TEV cleavage site, or with the pET14b vector encoding MnmG preceded by an N-terminal His-tag and TEV cleavage site. For both proteins, the first purification step consisted of Ni^2+^-NTA immobilized metal affinity chromatography using a His-trap FF column (5mL, Cytiva). After this first Ni^2+^-NTA purification step MnmE was subjected to an overnight alkaline phosphate treatment to hydrolyze the bound nucleotides, followed by a second Ni^2+^-NTA IMAC step to remove the alkaline phosphate. As a final purification step for MnmE and MnmG, the proteins were loaded on a Superdex200 size exclusion chromatography column and eluted using 20mM HEPES pH7.5, 150mM NaCl, 5mM MgCl_2_, 2mM DTT.

### Cryo-EM sample preparation and data acquisition

To obtain the MnmE-MnmG complex, 100 µM MnmE and 50 µM MnmG were mixed in the presence of 1 mM GppNHp and 1 mM FAD in a buffer containing 20 mM HEPES pH 7.5, 150 mM KCl, 5 mM MgCl_2_, 2 mM DTT. Subsequently, the mixture was loaded on a Superdex200 10/300 column and eluted with the same buffer supplemented with 100 µM GppNHp. The peak fractions corresponding to the α_4_β_2_ complex were pooled and immediately supplemented with 100 µM FAD. The molecular mass distribution of this fraction at a concentration of 100 nM was determined using mass photometry on a Refeyn OneMP instrument.

Subsequently, 3 µL samples at a protein concentration of 0.4-0.6 mg/ml were loaded on Quantifoil 1.2/1.3 300-mesh copper Holey grids priorly glow discharged for 40-60 s using an ELMO glow Discharger (Corduan Technologies). The grids were blotted for 2.8-3.6 seconds using Whatman No. 2 paper and plunge frozen in liquid ethane with a Gatan Cryoplunge3 and stored in liquid nitrogen until use. Two single-particle cryo-EM datasets were collected using a 300kV JEOL Cryo ARM 300 transmission electron microscope, equipped with an omega energy filter using a slit width of 20 eV. Images were captured with a K3 detector (Gatan) operating in correlative-double sampling (CDS) mode using SerialEM v3.0.8. Each micrograph consists of 60 frames with total exposure time of 3.036 s and a total dose of 62 e^-^/ Å^2^. Data were collected at a nominal magnification of 60,000x, corresponding to a pixel size of 0.76 Å. A total of 3,384 movies were collected for the first dataset and 19,370 movies for the second dataset (**Fig. S1**). Data collection statistics are detailed in **Table S1**.

### Image processing

Data processing, including gain normalization, motion correction and dose weighting, was performed using Relion 3.1^3^ and UCSF MotionCor2^4^. For the first dataset, images were curated with the in-house script BXEMDALYZER (Shkumatov et al., in preparation), while for the second dataset cryoSPARC v4.5 was used for curating images and analysis. The movies or motion-corrected micrographs were imported into cryoSPARC v4.5^5^ and CTF was calculated using Patch CTF.

For the first dataset, blob picking on a small subset of 50 images was performed, followed by 2D classification and manual curation to remove bad particles. The remaining particle stack was used to train a model for Topaz [39]. Topaz extraction was performed on Topaz denoised images, which resulted in a total of 164,809 picked particles. Several rounds of 2D classification and selection resulted in a particle stack of 122,647 particles used for *ab initio* modelling and subsequent heterogeneous refinement. Three classes were discerned, corresponding to the α_4_β_2_ complex, the α_2_β_2_ complex, and a class containing low quality smaller particles that possibly correspond to MnmE (α_2_) or MnmG (β_2_). This resulted in a total of 41,635 particles for the α_4_β_2_ complex class, 45,868 particles for the α_2_β_2_ class and 35,144 particles for the remaining class. Particles from the first two classes were kept for further refinement. To pick particles from the second dataset, templates obtained from 2D classification from the first dataset (corresponding to the α_2_β_2_ and α_4_β_2_ complexes) were used for template picking on cryoSPARC, which produced 1,947,753 particles. Several rounds of 2D classification and selection resulted in 355,130 particles used for *ab initio* reconstruction and heterogenous refinement into three classes. However, since these classes only corresponded to the α_2_β_2_ complex and to smaller particles corresponding to MnmG dimers (β_2_), only the particles from the former class were kept for further analysis and merged with the particles from dataset 1. All particles were aligned to place the center of the box at the center of the MnmG dimer, allowing for a more accurate 3D classification and separation into the α4β2 and α2β2 complexes, by using a mask on the second MnmE dimer. Homogeneous refinements yielded 116,023 particles for the α_2_β_2_ complex and 53 447 particles for the α_4_β_2_ complex. Before further refinement, particles were reextracted with a box size of 512 pixels for the α_2_β_2_ complex and 672 pixels for the α_4_β_2_ complex. Homogeneous refinement, followed by NU-refinement resulted in a 3.3 Å map reconstruction for the α_2_β_2_ complex using C1 symmetry, and a 4.0 Å map for the α_4_β_2_ complex using C2 symmetry. For the α_4_β_2_ complex one focused map was calculated centered on the MnmG dimer using a symmetry expanded particle stack. We obtained resolutions of 4.25 Å for the MnmG focused map, 4.48 Å for the MnmE A-B dimer and 4.60 Å for the MnmE E-F dimer (**Table S1, Fig. S1**).

### Model building and refinement

Prior to model building, the interpretability of the map of the α_2_β_2_ complex was improved by density modification in resolve_cryo_EM [40] within the PHENIX suit (version 1.21) [41]. Initial models of *E. coli* MnmG and *E. coli* MnmE were generated using Alphafold2 [42]. The resulting MnmG model was placed in the density map using rigid body fitting in UCSF ChimeraX 1.7 [43]. For model building of MnmE, first, the X-ray crystal structure of the GDP.AlF_4_-bound G domain dimer of MnmE (PDB 2gj8, [30]) was placed in the EM map. The MnmE α-helical domains and N-terminal domains were subsequently manually placed one by one in the map. Due to large conformational changes compared to the AlphaFold2 model in the subunit B of MnmE, flexible fitting was performed in Coot 0.9.8 before further refinement [44]. Real-Space refinement was performed in PHENIX using phenix.real_space_refine, applying secondary structure and Ramachandran restraints, followed by manual improvements in Coot. Subsequently, the chains of the final α_2_β_2_ model were placed into the general map for the α_4_β_2_ complex using rigid body fitting in ChimeraX. Focused maps were used to facilitate interpretation and model building of the corresponding regions. After refinement, these partial models were merged into the full model, and further refinement cycles followed by model validation were performed (**Table S1**).

### Cα distance matrix analysis

In the model of the α_2_β_2_ complex Cα distances were calculated for all MnmE subunits to generate Cα distance matrices. These matrices were compared between subunits A and B of MnmE to identify structurally conserved domains between the conformations.

### Unbiased molecular dynamics simulations

The structures of the MnmE subunits in their two conformations were extracted from the model of the α_2_β_2_ complex. Both subunits A and B were solvated using TIP3P water in a rhombic dodecahedron simulation box leaving at least 12 Å between the protein and the edge of the simulation box; Na^+^ and Cl^−^ ions were placed at a concentration of 100 mM neutralizing any net charge of the protein. For the protein the CHARMM36m force field was used [45]. The simulations were carried out with GROMACS 2023.1 [46]. The leap-frog algorithm was used with an integration time step of 2 fs. Non-bonded interactions were treated with a Verlet list cutoff scheme with a cutoff of 1.2 nm. Lennard-Jones potentials are smoothly switched to zero (Potential-switch) between 1.0 nm and 1.2 nm. The particle mesh Ewald method was used to treat long range electrostatic interactions with a grid spacing of 0.12 nm. The LINCS algorithm was used to constrain bonds with hydrogen atoms. Each molecular system was minimized for 10,000 steps of steepest descend.

Temperature equilibration at 298 K was performed over 50 ps of simulation under NVT ensemble using the v-rescale thermostat with separate heat bath couplings for solute and solvent [47]. Pressure equilibration at 1 bar was done over 500 ps of simulation under NPT ensemble using the c-rescale pressure coupling. During both equilibration simulations positional restraints of 1000 kcal mol^−1^ nm^-2^ were applied to the proteins. The production simulations were performed for 500 ns under NPT ensemble without positional restraints. The v-rescale thermostat and c-rescale barostat were used for temperature and pressure coupling, respectively. Conformational changes throughout the simulations were analyzed using GROMACS’ *gmx rms* module using both the native states of subunit A and subunit B as references.

### Biased molecular dynamics simulations of the MnmE conformational transition

To find a potential transitioning pathway between the two MnmE subunit conformations, an iterative procedure was applied. Separate biased simulations are stated from each endpoint of the transition. Prior to each simulation step, the structures are superimposed and vectors between corresponding Cα atoms are calculated. Along these vectors a harmonic potential is applied that biases the simulation towards the other subunit’s conformation. The effective force acting on each Cα is adjusted by moving the center of the harmonic potential along the Cα-Cα vector. Applying this bias, a short simulation of 500 ns is run from both endpoints. After this step, the endpoints of both simulations are extracted, superimposed and Cα vectors are calculated to define biases for the subsequent step of the simulation. This process is repeated iteratively until the remaining RMSD between the endpoints is less than a 1.2 Å.

### Analysis of small angle X-ray scattering (SAXS) data

Experimental SAXS data of MnmE bound to GDP*AlF_4_, MnmE bound to GppNHp and of the MnmE-MnmG complexes in their α_2_β_2_ and α_4_β_2_ states were previously obtained as described in Fislage *et al.*, 2014 [38]. These experimental SAXS data were compared with the theoretical SAXS curves corresponding to the models of the α_2_β_2_ and α_4_β_2_ complexes obtained using cryo-EM in this study, or to the asymmetric MnmE dimer extracted from the α_2_β_2_ cryo-EM model, using CRYSOL [48] from the ATSAS software package [49]. CRYSOL was run using the number of spherical harmonics set to 50 to improve the precision of the theoretical scattering profiles, and with a constant subtraction to compensate for a potential improper background subtraction of the SAXS data.

## Results

### Cryo-EM structures of the MnmE-MnmG complex show the existence of α_2_β_2_ and α_4_β_2_ states

Previous SAXS modelling of the complex formed between the MnmE (α_2_) and MnmG (β_2_) homodimers, suggested the presence of an L-shaped oligomer with α_2_β_2_ stoichiometry when MnmE is in its nucleotide-free state or bound to GDP. Upon binding of GTP to the G domains of MnmE, formation of a higher oligomeric state with α_4_β_2_ stoichiometry was observed, which returns to an α_2_β_2_ complex upon GTP hydrolysis [38]. Here, we prepared the MnmE-MnmG complex (MnmEG) in presence of potassium, the non-hydrolysable GTP analogue GppNHp and the MnmG cofactor FAD. This complex was purified by SEC, and the mass was determined using mass photometry immediately prior to applying the sample on cryo-EM grids. While mass photometry indicated some heterogeneity in the mass distribution within the purified sample, more than 70% of the population attains a molecular mass close to the 348 kDa expected for the α_4_β_2_ complex (**Fig. S2)**.

After optimization of the grid conditions, two datasets were collected on the JEOL Cryo-ARM 300 electron microscope. Within the first dataset, *ab initio* modelling and subsequent hetero-refinement and 3D classification led to the identification of two nearly equally populated classes: one corresponding to the very elongated α_4_β_2_ complex and one corresponding to the α_2_β_2_ complex (**Fig. 1B Fig. S1**). Although the complex that was used to collect the second dataset was purified in the same way, the main particle species observed here corresponded to the α_2_β_2_ complex. This suggests that the α_4_β_2_ complex, which is the main species prior to grid preparation according to mass photometry, partially dissembles into the α_2_β_2_ complex upon grid preparation and plunge freezing (**Fig. S3**). Further processing to obtain high quality reconstructions of both complexes was done by merging particle stacks of both datasets and by optimizing the alignment during 2D and 3D classification and by particle re-extraction aligning the middle of the MnmG dimer to the box center. A thorough 3D classification by applying a mask on the second MnmE molecule allowed us to better distinguish between the α_4_β_2_ and α_2_β_2_ classes, yielding 53,447 particles for the α_4_β_2_ complex and 116,023 particles for the α_2_β_2_ complex. Further refinement resulted in global resolution estimates of 4.0 Å for the α_4_β_2_ complex and 3.3 Å for the α_2_β_2_ complex (**Fig. S1, Table S1**). In addition, the EM map of the α_4_β_2_ complex showed clear C2 symmetry, and an MnmG focused map was obtained by performing local refinement using C2 symmetry-expanded particle stacks and applying masks on the MnmG dimer.

To build the α_2_β_2_ model, AlphaFold2 models for *E. coli* MnmG and MnmE were first fitted into the EM maps [42], followed by manual building in COOT and refinement in Phenix, and placement of the ligands FAD and GppNHp if density was present. For MnmG only minor conformational changes were apparent allowing rather straightforward placement of the Alphafold2 model, with only some adjustments including the deletion of a large part of the helical domain in one of the subunits (subunit D). In contrast, the MnmE subunit that interacts with MnmG (subunit B) is subject to very drastic conformational changes compared to the Alphafold2 model, necessitating to fit each (sub)domain of MnmE individually in the map (**Fig. 1C**). Reassuringly, the resulting model of the α_2_β_2_ complex fits well with our previously collected SAXS data of this complex in solution, with a χ^2^ value of 2.8 (**Fig. S4A**) [38]. The resulting models and the observed conformational changes will be described in detail in the subsequent paragraphs.

To build the α_4_β_2_ complex, first the MnmE asymmetric dimer was taken from the α_2_β_2_ model and fitted in an identical fashion in the region of the α_4_β_2_ map corresponding to the MnmE A and B subunits (with MnmE subunit B interacting with MnmG subunit C). In the same way, MnmE was placed in the region of the map corresponding to the MnmE E and F subunits (with MnmE subunit E interacting with MnmG subunit D). To place MnmG in the α_4_β_2_ complex, a locally refined map of the MnmG dimer was used. Chain C fitted in an identical way as in the α_2_β_2_ complex. However, clear differences were observed in MnmG subunit D compared to the α_2_β_2_ complex, where density is now also observed for the helical domain. As such, the locally refined map could convincingly be fit with a symmetrical MnmG model. Finally, a general model for the α_4_β_2_ complex was built by combining the partial MnmG model from the local map with the two MnmE dimers and fitting these into the full density map for the α_4_β_2_ complex (**Fig. 1D, Table S1**). Also for the resulting α_4_β_2_ complex we find a good fit with the SAXS data of the corresponding complex in solution, with a χ^2^ value of 2.6 (**Fig. S4B**) [38].

Considering the higher resolution of the α_2_β_2_ complex compared to the α_4_β_2_ complex, the former will be used for most of the subsequent analyses and discussions, unless stated otherwise.

### MnmG gets structured and binds FAD upon interaction with MnmE

The structure of the α_2_β_2_ complex shows an MnmG dimer (β_2_), with each subunit consisting of the typical Rossmann-fold FAD-binding domain (residues 1-42, 100-160, 175-192, 349-397), interrupted by 2 insertion domains (insertion domain 1: 43-99; insertion domain 2: 161-174, 193-348) and a C-terminal α-helical domain (398-629) [32]. However, while previous crystal structures of MnmG from *E. coli* (EcMnmE), *Chlorobaculum tepidum* (CtMnmG) and *Aquifex aeolicus* (AaMnmG) present MnmG as a symmetric homodimer, the subunits (C and D) of MnmG within the α_2_β_2_ complex are not identical, due to the structuring of flexible regions in subunit C upon interaction with MnmE (**Fig. 2**) [32–34].

**Figure 2:**
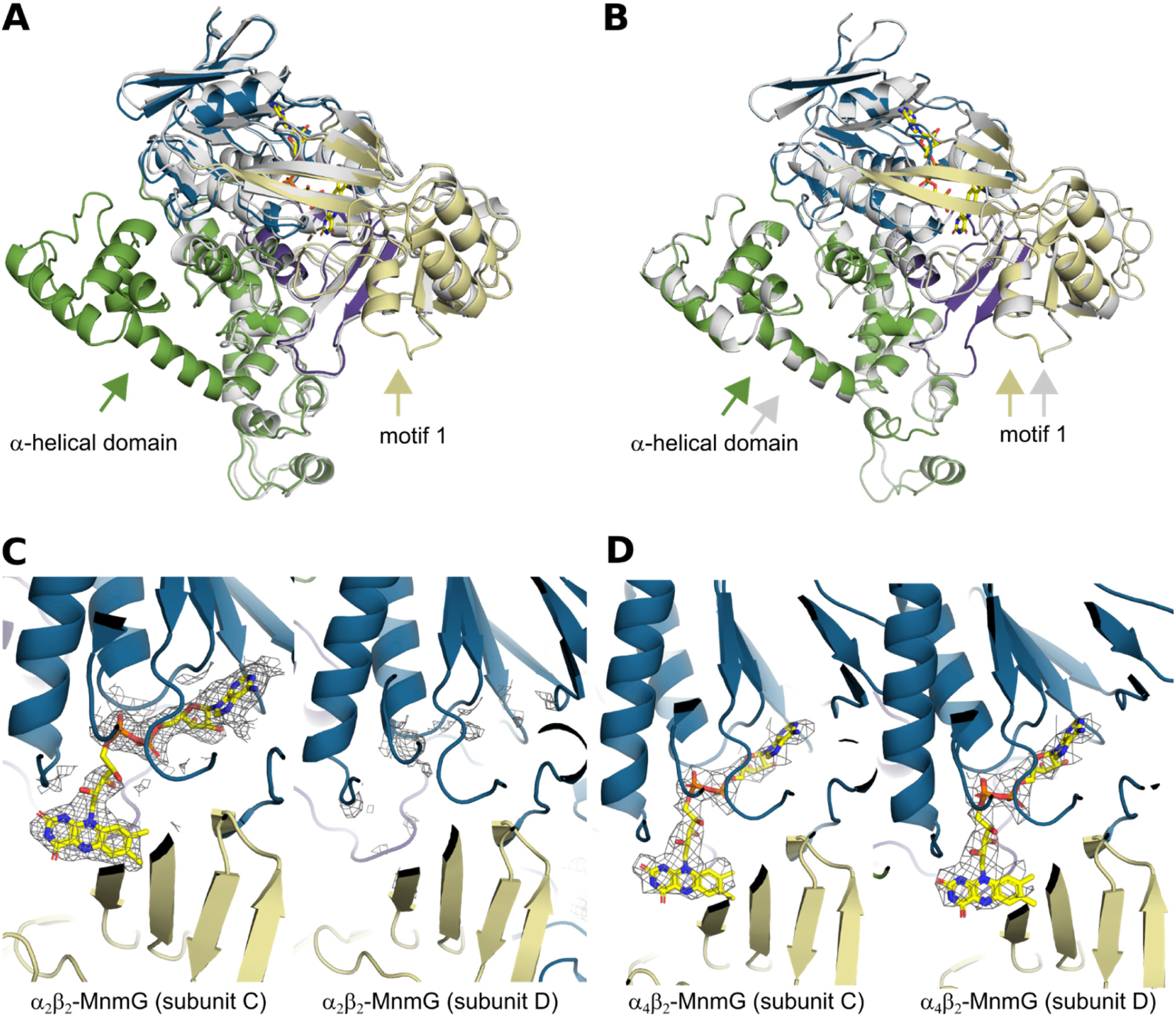
Structural changes and FAD binding of MnmG upon interaction with MnmE. **(A)** Superposition of the C and D subunits of the MnmG dimer within the α_2_β_2_ MnmEG complex. In the α_2_β_2_ complex the interaction with MnmE is made by the MnmG subunit C. Subunit D is shown in grey, while subunit C is colored according to its domains, with the FAD domain, insertion domains 1 and 2 and the helical domain colored in blue, dark purple, beige and green, respectively. The structuring of the α-helical domain and MnmG-specific motif 1 in subunit C, upon interaction with MnmE, is indicated with arrows. **(B)** Superposition of the C and D subunits of the MnmG dimer within the α_4_β_2_ MnmEG complex, in which both MnmG subunits interact with MnmE. Correspondingly, the α-helical domain and MnmG-specific motif 1 is structured in both subunits as indicated by arrows. **(C)** and **(D)** Cryo-EM density map, shown as a grey mesh, around the FAD binding site of subunit C and D of MnmG within the α_2_β_2_ (**C**) and α_4_β_2_ (**D**) MnmEG complexes. The density in both subunits is shown at the same contour level.

A first noticeable difference between the C and D subunits of MnmG concerns their C-terminal α-helical domains. In previous crystal structures of EcMnmG a large part of this domain (residues 554-629) could not be modelled due to flexibility, while in more rigid MnmG orthologues from thermophiles these regions were folded in well-structured α-helical arms [32–34]. Similar to the EcMnmG crystal structures, this region shows no density in the D subunit of the α_2_β_2_ complex (**Fig. 1C, Fig2A**). In contrast, in the C subunit the α-helical domain is involved in intimate interactions with MnmE leading to a disorder-order transition where a fold is attained that is very similar to the one observed in the CtMnmG and AaMnmG crystal structures. Correspondingly, in the α_4_β_2_ complex both α-helical domains of the MnmG dimer are structured upon interaction with MnmE, and the symmetry in MnmG is restored (**Fig. 1D, Fig2B, Fig. S5**).

A second important region of asymmetry and conformational change entails the so-called MnmG-specific motif 1, which is located in insertion domain 2 and was previously hypothesized to be involved in binding of the substrate NADH [32]. This motif (residues 273-287) is part of a larger unstructured peptide region (residues 255-292) in MnmG crystal structures, as well as in the D subunit of the α_2_β_2_ complex. However, in subunit C of the complex most of this region gets ordered and is implicated in important interactions with the C-terminal helix of MnmE (**Fig. 2A&B**).

Interestingly, we also observe very clear density in the EM map corresponding to a bound FAD molecule in subunit C of the MnmG dimer, while no such density is present in subunit D (**Fig. 2C**). EcMnmG binds FAD with relatively low affinity (K_D_ ≍ 3 µM), and, correspondingly, loses FAD during size exclusion chromatography [34]. Although 100 µM of FAD was added to the MnmE-MnmG complex prior to cryo-plunging, the structure clearly shows asymmetric binding of FAD in only one of the two MnmG subunits. Closer inspection of the structure shows that MnmE binds with its α-helical domain on top of the deep FAD binding pocket of MnmG, thereby partially closing off this pocket. Moreover, within the complex the side chain amino group of the C-terminal lysine residue (Lys454) of MnmE is located within interaction distance of the C(4)=O carbonyl group of the isoalloxazine moiety of FAD. This strongly suggests that MnmE is involved and supports the binding of FAD to MnmG. In agreement with this observation, we find density for bound FAD molecules in both MnmG subunits of the α_4_β_2_ complex (**Fig. 2D**).

### MnmE undergoes large scale conformational changes and adopts an asymmetrical arrangement in complex with MnmG

In contrast to all previously solved crystal structures that show MnmE as a symmetric homodimer in an “open” conformation with the G domains of each subunit facing but not contacting each other, the GppNHp-bound MnmE dimer (α_2_) within the α_2_β_2_ complex adopts a “closed” conformation, where the two G domains contact each other [27,28]. This agrees with fluorescence spectroscopy and EPR studies, which suggested an “open” to “close” transition of the MnmE G domains upon GTP binding and/or interaction with MnmG [18,28,31,50,51]. Although the local resolution around the G domains is rather low, the EM map agrees with the crystal structure of the GDP*AlF_4_-bound G domain dimer of EcMnmE (PDB code 2GJ8) [30].

However, in the α_2_β_2_ complex the MnmE dimer shows very pronounced differences in conformation between the subunit that is located at the periphery of the α_2_β_2_ complex (A subunit) and the subunit contacting MnmG (B subunit) (**Fig. 3**). The overall conformation and the domain arrangement of the A subunit resemble those of the crystal structures of MnmE, with an N-terminal 5,10-CH_2_-THF binding domain (a.a. 1-119) consisting of a five-stranded mixed β-sheet and 3 α-helices, a helical domain (120-215 and 377-454) consisting of 8 α-helices (Hα1 – Hα8), among which the long helices Hα3-Hα5-Hα6-Hα7 form a 4-helix bundle, and a G domain that is inserted between Hα5 and Hα6 of the helical domain (**Fig. 3A, Fig. S6)** [17]. The C-terminal helix of the helical domain (Hα8) of MnmE subunit A interacts with the N-terminal domain, and the very last loop residues that contain the highly conserved FC(V/I/L)GK motif are inserted into the 5,10-CH_2_-THF binding pocket of the N-terminal domain. In contrast, MnmE subunit B, that interacts with subunit C of MnmG, undergoes a number of very drastic and unanticipated conformational changes. Indeed, superposition of the N-terminal domains of the A and B subunit of MnmE shows a very large rotational movement of the helical domain of subunit B with respect to the N-terminal domain of nearly 112° (measured as the angle between the Cα atoms of the fixed residue N118 on the N-terminal domains and residues A113 at the “tip” of the helical domain of subunit A and B) (**Fig. 3B**). The main driver of this conformational change seems to be a large rearrangement in the region connecting the N-terminal domain to the helical domain (a.a. 120-150). Similar to all currently solved crystal structures of MnmE, in the A subunit of MnmE, this region is arranged in two short and nearly anti-parallel oriented α-helices (Hα1 consisting of a.a. L123-A135 and Hα2 consisting of a.a. E138-Q149) connected by a short loop. However, in the B subunit that directly interacts with MnmG, these two helices rearrange into one long helix (a.a. A126-Q149), accompanied by the translocation of the entire helical domain (**Fig. 3C**). Considering the crucial role of this helix in inducing the observed conformational changes, we called this the “hinge helix”. Interestingly, two residues that are located within the hinge helix (E127 and A130) correspond to pathogenic disease mutations linked to hypertrophic or dilated cardiomyopathy in human GTPBP3 (E159V and A162P) [22], underscoring the importance of the hinge helix in maintaining the physiological function of MnmE-MnmG / GTPBP3-MTO1 (**Fig. 3C**).

**Figure 3:**
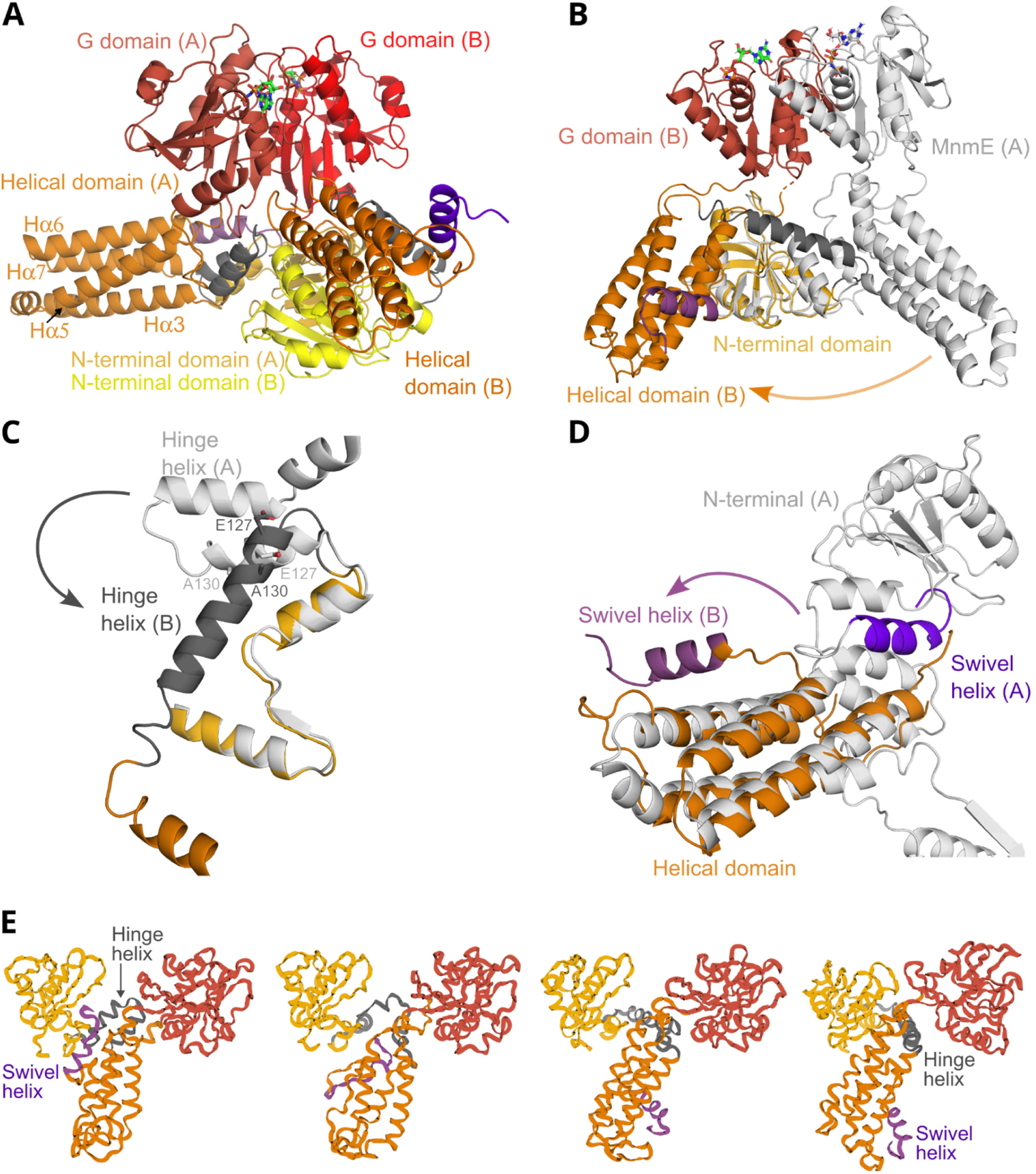
Conformational changes in the MnmE dimer upon interaction with MnmG within the α_2_β_2_ MnmEG complex. **(A)** The asymmetric MnmE dimer as found within the α_2_β_2_ complex. In the α_2_β_2_ complex the interaction with MnmG is made by the MnmE subunit B. The domains belonging to the A and B subunits are indicated in different color shades, with the N-terminal domain, helical domain and G domain, shown in yellow, orange and red, respectively. The hinge helix and swivel helix are shown in dark grey and purple clarity, subunit A is shown in light grey, while subunit B is colored according to its domains as in panel (A). This superposition shows a very large rotational movement of the helical domains with respect to the N-terminal domains by about 112°. **(C)** Superposition of the A and B subunits of MnmE similar to (B), zooming in on the large conformational change in the hinge helix. Two shorter helices (L123-135 and E138-Q149) in subunit A rearrange into one long helix in subunit B. The positions of E127 and A130, which correspond to the location of disease-associated mutations in GTPBP3, are also indicated. **(D)** Superposition of the A and B subunits of MnmE similar to (B), showing the large change in the position of the swivel helix. **(E)** Snapshots along the biased molecular dynamics simulation of the pathway from the MnmE “subunit A conformation” toward the “subunit B conformation”. The color code used is similar to panel (A). The trajectory seems to be driven by a “twist” in the helical domain facilitated by partial unfolding of the swivel helix. The swivel helix then relocates from its position in the N-terminal domain to the helical domain concomitant with the rearrangements of the hinge helix.

Further comparison of the helical domains of the A and B subunits of MnmE reveals additional changes. A first important difference concerns the C-terminal helix (Hα8) and the adjacent FCIGK motif. While in the A subunit this region is inserted into the 5,10-CH_2_-THF -binding pocket of the N-terminal domain, in the B subunit it has undergone a huge rearrangement and now forms direct interactions with the FAD-binding region of MnmG (**Fig. 3D**). We therefore named this helix the “swivel helix” and we will discuss its functional implications in more detail later. Apart from this very marked difference, also some other less pronounced conformational changes can be observed in the helical domain of the B subunit, especially in the region connecting helices Hα3 to Hα5 (a.a. 173-188).

In the α_4_β_2_ complex similar conformational changes, albeit at lower resolution, can be observed between the subunits (A *vs.* B or E *vs*. F) of both MnmE molecules bound at either side of MnmG. These asymmetrical conformational changes in the α_2_β_2_ and α_4_β_2_ complexes could in principle be triggered by either binding of GppNHp, by binding to MnmG, or require a combination of both. Indeed, while previous fluorescence spectroscopy and EPR studies showed that binding of the non-hydrolysable GTP analogue GppNHp or the transition state analogue GDP*AlF_4_ induce the closing of the MnmE G domains, it is unclear whether this also induces the asymmetrical large-scale conformational changes observed here [28,31,50,51]. To address this question, we compared the theoretical SAXS curve calculated from the structure of the MnmE dimer extracted from the α_2_β_2_ complex to the previously obtained experimental solution SAXS curves of MnmE bound to either GDP*AlF_4_ or GppNHp (**Fig. S4C & S4D**) [38]. This analysis shows that there are very significant differences in the overall shape of GDP*AlF_4_- or GppNHp-bound MnmE in solution and the conformation of MnmE in complex with MnmG, thus clearly indicating that the asymmetrical conformational changes observed in MnmE are at least partially triggered by the binding to MnmG, and hence are unique to the MnmE-MnmG complex.

### The conformational changes in MnmE shape the MnmE-MnmG interface

The commonly buried accessible surface area between MnmE and MnmG in the α_2_β_2_ complex amounts to 2134 Å^2^ (calculated via the PISA tool on PDBe) [52]. Closer inspection allows to distinguish two main interaction surfaces (**Fig. 4**). The largest surface (interaction surface 1) is formed by interactions between the displaced helical domain of MnmE on the one hand, and residues of the insertion domain 1, insertion domain 2 and helical domain of MnmG on the other hand (**Fig. 4A**). A first patch within this region consists of residues from the loop connecting Hα3 to Hα5 from the MnmE helical domain (a.a. E171-G187), which interact with residues of insertion domain 1 (a.a. G53-G55 and R82-V92), insertion domain 2 (a.a. R290-F296) and the helical domain (a.a. R427-S431) of MnmG. Prior data already showed the importance of several of these involved MnmE residues for binding to MnmG [31,38]. Moreover, we previously showed that mutation of MnmE residue D175, which interacts with S431 of MnmG in our structure, impairs the tRNA modification reaction [31]. Additionally, MnmG residue G55 within this interaction region corresponds to the site of a cardiomyopathy-associated mutation in human MTO1 (G85R) [53,54]. A second patch within interaction surface 1 consists of residues from the loop connecting the other two helices of the MnmE 4-helix bundle (Hα6 and Hα7; a.a. 405-425) that interact with residues of the MnmG insertion domain 1 (a.a. S87-A91) and helical domain (a.a. E434-L437, P516-E517, Y548-Q555). Our previous mutagenesis studies also showed a role of MnmG residues Y551 and R554 in the interaction with MnmE [38]. Moreover, the corresponding regions in GTPBP3 and MTO1 seem to be affected by pathogenic mutations, including the E459K mutation in GTPBP3 (corresponding to E421 in MnmE) and R473C in MTO1 (corresponding to R436 in MnmG) [22,55]. A third patch of interaction surface 1 is dominated by the MnmE swivel helix. Upon interaction with MnmG the swivel helix is displaced from its original position close to the N-terminal domain of MnmE toward the FAD-binding pocket of MnmG. In this position residues from the swivel helix form multiple interactions with residues from the insertion domains 1 and 2 of MnmG. Within MnmG insertion domain 2, specifically residues that constitute the highly conserved MnmG-specific motif 1 are involved. This MnmG-specific motif seems to be a hot-spot for disease mutations in MTO1, with the R313Q and K321N mutations (corresponding to EcMnmG residues R275 and K283, respectively) found implicated in hypertrophic cardiomyopathy [53,55,56].

**Figure 4:**
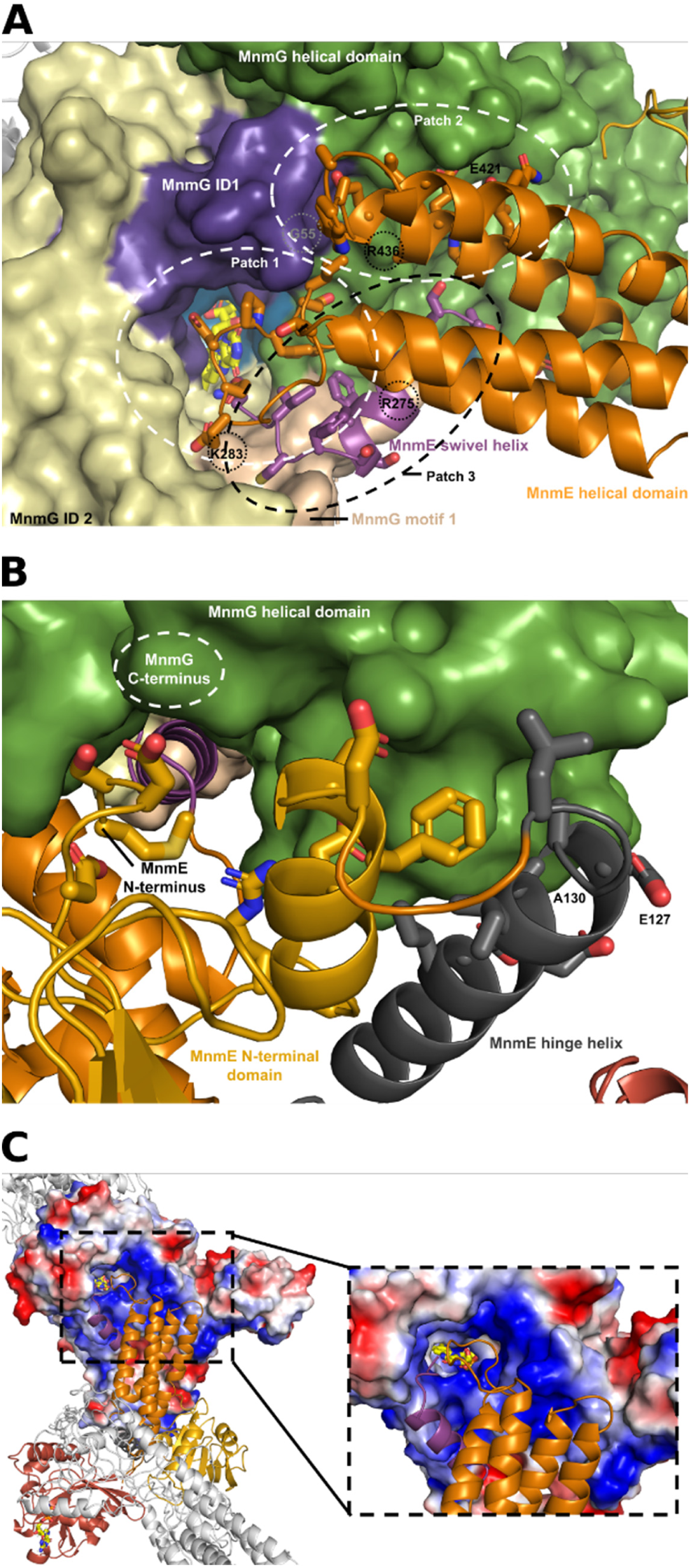
The MnmE-MnmG interaction surface. (**A**) Close-up view on MnmE-MnmG interaction surface 1. Subunit C of MnmG is shown in surface representation with the FAD domain, insertion domains 1 and 2 (ID1and 2) and the helical domain colored in blue, dark purple, beige and green, respectively. The MnmG-specific motif 1, which is part of ID2, is represented in the color wheat. The FAD molecule is shown in stick representation with the C-atoms colored yellow. MnmE is shown in cartoon representation with the N-terminal domain and helical domain in yellow and orange, respectively. The swivel helix is represented in light purple. The three major patches that contribute to the interaction, as described in the text, are indicated. Residues of MnmE that make interactions with MnmG are shown as sticks. Residues of MnmE and MnmG that correspond to the location of disease-associated mutations in GTPBP3 and MTO1 are also indicated. (**B**) Close-up view on MnmE-MnmG interaction surface 2. The mode of representation and colors are the same as in (A). The MnmE hinge helix is indicated in dark grey. Residues of MnmE that correspond to the location of disease-associated mutations in GTPBP3 and MTO1 are indicated. (**C**) MnmE binds on a positively charged surface patch surrounding the FAD-binding pocket of MnmG. The electrostatic potential surface around the C subunit of MnmG is shown, together with the interacting B subunit of MnmE in cartoon representation. The color code used for MnmE is the same as in (A) and (B). The non-interacting A and D subunits of MnmE and MnmG, respectively, are also shown in grey cartoon representation. The right panel shows a zoom-in around the FAD-binding pocket of MnmG, with FAD shown in stick representation.

A second interaction surface (interaction surface 2) is formed by interactions of the N-terminal domain and the elongated hinge helix of MnmE with the C-terminal part of the helical domain of MnmG (**Fig. 4B**). In particular, this interface is dominated by a 3-helix bundle formed by the last α-helix of the N-terminal domain of MnmE (a.a. R107-N119), the hinge helix of MnmE (a.a. Q125-R142) and the helix formed by residues T608-Q622 of MnmG. An additional, less extensive, patch of interaction surface 2 is formed by the N-terminal residues of MnmE (a.a. M1-D5) with α-helix K547-N567 of MnmG. Residues of GTPBP3 located on interaction surface 2 were also identified as disease mutations, including E159V and A162P (corresponding to MnmE E127 and A130) [22].

### Is the MnmE swivel helix acting as a relay of reaction intermediates?

Our current structures also provide a plausible mechanism for the transfer of a methylene group from the 5,10-CH_2_-THF donor to the FADH2 acceptor to form a flavin iminium intermediate, as suggested in the most recently proposed reaction mechanisms [36,37]. In this scenario the swivel helix could serve as a relay system to transfer the methylene group from the 5,10-CH_2_-THF molecule bound to MnmE to the FADH_2_ molecule bound to MnmG. In previously solved crystal structures of MnmE, as well as in subunit A of MnmE in the current α_2_β_2_ complex, the swivel helix and the adjacent C-terminal coil region harboring the highly conserved ^450^FCIGK^454^ motif are protruding into the 5,10-CH_2_-THF-binding pocket located in the N-terminal domain of MnmE [27,28]. In this position the swivel helix is held in place via a 3-helix bundle involving the last α-helix of the N-terminal domain (E110-N118) and the first α-helix of the helical domain (L123-I133) (**Fig. 5A**). The latter helix constitutes the N-terminal half of the hinge helix. As such, there is a direct interaction between the swivel and hinge helices, constituting a mechanism to propagate the observed large conformational changes from one helix to the other upon interaction with MnmG. Interestingly, while the side chain of the highly conserved C-terminal K454 residue points away from the 5,10-CH_2_-THF-binding pocket in the A subunit of the α_2_β_2_ complex, previously solved crystal structures of MnmE bound to 5-formyl-THF show this side chain pointing toward the cofactor, suggesting a THF-induced conformational change. In the crystal structure of *Thermotoga maritima* MnmE bound to 5-formyl-THF (TmMnmE, PDB 1xzq) the amino group of K450 (K454 in EcMnmE) is located at 2.6 Å from the 5-formyl group, and K450 is further surrounded by Y71, R20, V126 and I130 (F73, R23, I129 and I133 in EcMnmE), and by Ala15 and His82 of the adjacent subunit (G18 and H84 in EcMnmE) (**Fig. 5A**) [27]. This environment of either non-polar or positively charged residues could tune the p*K*_a_ of K454 such that it is present in its neutral form. In this state it could perform a nucleophilic attack on the iminium tautomer of 5,10-CH_2_-THF, resulting in a transfer of the methylene group to the amino group of K454.

**Figure 5:**
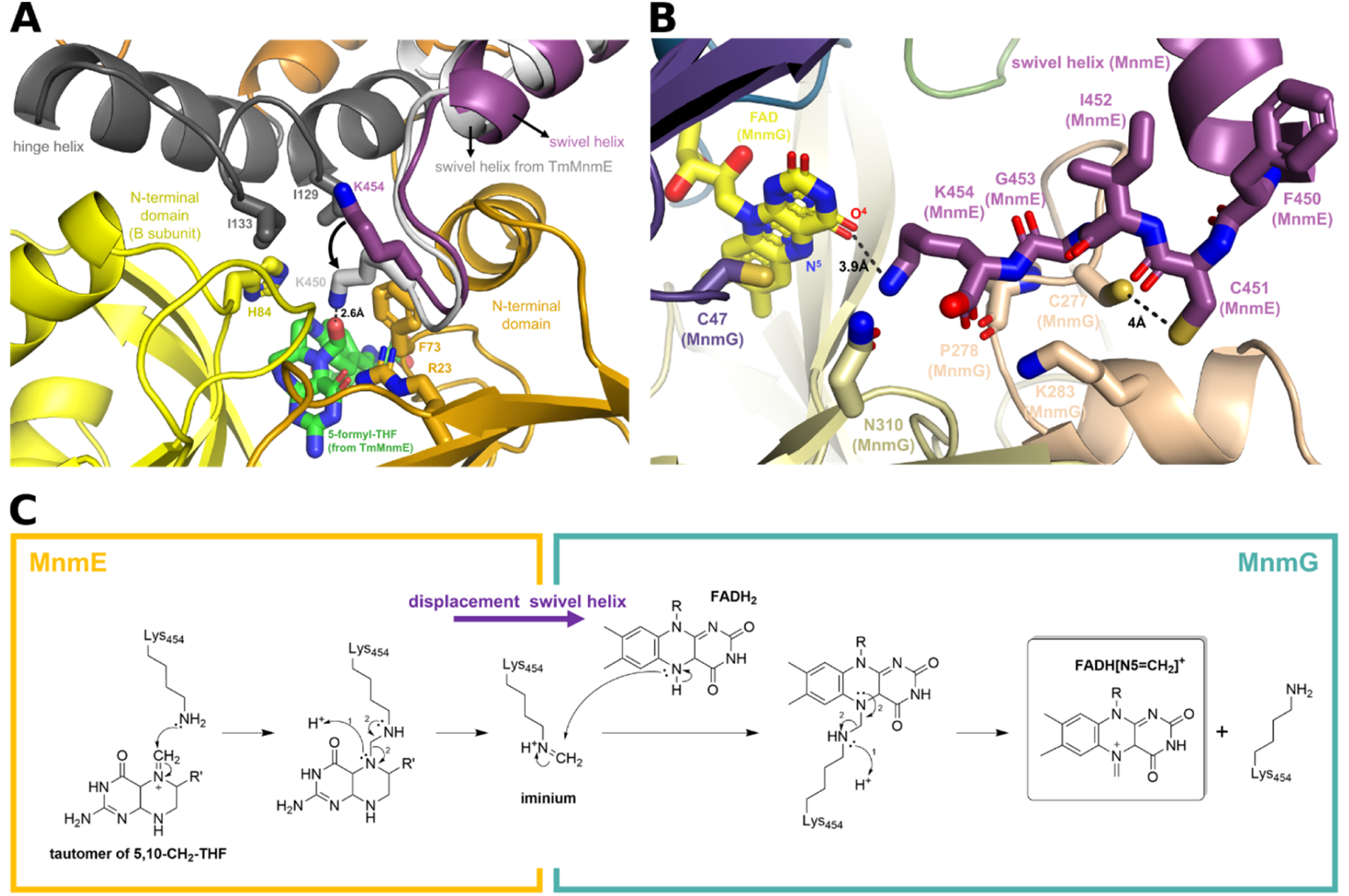
Molecular interactions of the swivel helix and the C-terminal ^450^FCIGK^454^ motif in the A and B subunits of MnmE, and potential implications for the tRNA modification reaction. **(A)** Position of the swivel helix in the 5,10-CH_2_-THF binding pocket of the A subunit of MnmE. The N-terminal and helical domains of the subunit A of MnmE are shown in dark yellow and orange, respectively, while the adjacent N-terminal domain of subunit B of MnmE is shown in light yellow. The A subunit hinge and swivel helices are shown in dark grey and purple respectively. The corresponding swivel helix of the superposed crystal structure of MnmE from *T. maritima* (TmMnmE, PDB 1XZQ) is shown in light grey, together with the bound 5-formyl-THF molecule (in green). An arrow indicates the suggested conformational change of K454 upon binding of a THF derivative. **(B)** Position of the MnmE B subunit swivel helix within the FAD binding pocket of the C subunit of MnmG. The swivel helix is shown in purple, while the MnmG insertion domain 1, insertion domain 2 and MnmG-specific motif 1 are colored in dark purple, beige and wheat, respectively. The MnmG-bound FAD molecule is shown in yellow. K454 interacts with the O^4^ atom and is positioned close (5.8 Å) to the reactive N^5^ atom of FAD. The catalytically important cysteines: C451 of MnmE and C47 and C277 of MnmG are located in the same pocket, with MnmE C451 and MnmG C277 being in close proximity. **(C)** Hypothetical role of the C-terminal lysine residue (K454) of the swivel helix in the transfer of a methylene group from the 5,10-CH_2_-THF methyl donor in the active site of MnmE toward the FADH_2_ cofactor in the active site of MnmG.

The large conformational changes in the B subunit of MnmE in the α_2_β_2_ complex go hand in hand with a displacement of the swivel helix toward the entrance tunnel that leads to the MnmG-bound FAD. Here, all residues of the conserved ^450^FCIGK^454^ motif of MnmE make multiple polar and non-polar interactions with MnmG. Additionally, the terminal COO^-^ group of MnmE seems to be crucial for anchoring the swivel helix deep into the FAD binding pocket of MnmG, where it is located within interaction distance to the side chains of K283 and N310 and the main chain C=O of P278. In this position, the side chain of K454 is oriented toward FAD, at 3.9 Å from the O^4^ group of the isoalloxazine group (**Fig. 5B**). The distance between the amine group of K454 and the reactive N^5^ atom of FAD is 5.8 Å, sufficient to allow the presence of a methylene group covalently attached to K454. It is thus tempting to speculate that the observed large conformational changes in MnmE upon interaction with MnmG enable the translocation of a methylene group, via the K454 residue of the swivel helix, from the MnmE-bound 5,10-CH_2_-THF to the MnmG-bound FADH_2_, whereafter it can be further transferred to the tRNA wobble uridine (**Fig. 5C**).

Finally, it is also worth noting that the location of the swivel helix in the entrance tunnel leading toward FAD, places the sulfhydryl groups of the catalytically important C451 of MnmE and C277 of MnmG in close proximity (4 Å) and in an appropriate orientation to interact with each other (**Fig. 5B**). The exact functional implications of this observation remain unclear to us at this moment.

### Molecular dynamics simulations suggest a trajectory for the conformational changes in MnmE

To get a better understanding of the nature of the structural transition occurring between the conformations observed in the subunits A and B of MnmE within the α_2_β_2_ complex, we first calculated Cα distance matrices for both subunits. These matrices describe a protein’s conformation in internal coordinates and thus allow for an alignment-independent comparison of the conformations in subunits A and B (**Fig. S7**). This analysis showed that both the N-terminal domain and the GTP-binding domain move largely as rigid bodies during the conformational transition. Also, the helices of the 4-helix bundle of the helical domain (Hα3, Hα5, Hα6, Hα7) retain their relative positions towards each other. This implies that the large conformational change observed in MnmE is mostly driven by a few highly variable regions that allow a conformational rearrangement of otherwise rigid domains. These flexible regions are primarily the intersections between the domains including the “hinge helix” and the C-terminal “swivel helix”.

We next conducted MD simulations of the individual MnmE subunits A and B to investigate whether the conformational transition occurs spontaneously. Both conformations exhibited significant internal motion, with maximum displacements exceeding 10 Å (**Fig. S8**). These elevated values are partly due to the non-globular structure of MnmE which increases the sensitivity of the RMSD to motions of individual domains. However, it also suggests that MnmE is probably stabilized by inter-subunit interactions within the homodimer. Despite the substantial internal motion in both conformations, no spontaneous transition between them was observed. Each conformation deviated considerably from its initial structure during the simulation but did not converge on the other conformation. This finding is consistent with our SAXS data which indicated that interaction with MnmG might be required to facilitate the transition from the A to the B subunit conformation.

To identify the transition pathway between the two conformations, we resorted to biased molecular dynamics simulations. Defining suitable biasing potentials was challenging due to the complex domain rearrangements involved in the transition. Ultimately an iterative approach was required, where simulations from both endpoint conformations were run until they converged. Harmonic potentials were applied to each Cα atom, biasing their motion towards the position of the corresponding Cα position in the other conformation. After 30 iterations the subunits A and B converged with a final RMSD of 1.2 Å. Interestingly, rather than involving large-scale conformational rearrangements of entire domains, the transition was driven by a “twist” in the helical domain facilitated by partial unfolding of the protein’s C-terminus, including the swivel helix (**Fig. 3E& Supplementary movie 1**). This segment relocates from its position in the N-terminal domain (in subunit A) to the helical domain (in subunit B). The twist also induces the rearrangement and “straightening” of the two short helices of the hinge region (in subunit A) into one long hinge helix (in subunit B). In interpreting these molecular dynamics simulations, it is important to note, however, that the pathway described may not represent the lowest-energy transition. This approach is inherently biased toward finding a pathway that involves the least conformational rearrangements. In addition, in a physiological context, the presence of additional factors, like MnmG, may assist the transition.

## Discussion

After a low-resolution model of the MnmE-MnmG complex was proposed over 10 years ago based on SAXS data [38], we now present the first high resolution cryo-EM structures of this tRNA modifying enzyme complex. These structures confirm the presence of the asymmetric α_2_β_2_ and α_4_β_2_ conformational states. In the α_2_β_2_ complex the MnmE (α_2_) and MnmG (β_2_) dimers assemble alongside each other in an elongated heterotetramer leaving one MnmE and one MnmG subunit vacant on each side of the complex. In the α_4_β_2_ complex the vacant MnmG subunit is occupied by a second MnmE dimer, resulting in an even more elongated arrangement. This configuration results in two vacant MnmE subunits on each side of the α_4_β_2_ oligomer. An explanation for this intriguing asymmetry can be found in the asymmetric arrangement of the subunits of the MnmE dimer in the MnmEG complexes. Indeed, upon interaction with MnmG the interacting subunit of MnmE undergoes a large conformational change breaking the symmetry within the MnmE dimer. The formation of a corresponding symmetrical MnmE dimer, in which both subunits would adopt a conformation that allows interaction with MnmG, is highly unlikely since this would either induce severe steric clashes between the G domains (**Fig. S9A**) or require a completely different interaction interface between the N-terminal domains of the constitutive MnmE dimer (**Fig. S9B)**.

Closer inspection of the interacting MnmE and MnmG subunits within the α_2_β_2_ and α_4_β_2_ complexes reveals significant conformational changes in both protein partners. In MnmG the C-terminal α-helical domain and the loop region spanning the so-called MnmG-specific motif 1 undergo a disorder-to-order transition and fold into structural elements that take part in the interaction with MnmE. The conformational changes in the interacting subunit of MnmE are even more pronounced, and include a large rotational movement of the helical domain with respect to the N-terminal domain. We identified two key structural elements that drive these conformational changes in MnmE: the C-terminal “swivel helix” that contains the functionally important and conserved ^450^FCIGK^454^ motif, and the “hinge helix” that connects the N-terminal domain to the helical domain (**Fig. 6**). Upon binding to MnmG, the hinge helix rearranges from two short and nearly anti-parallel α-helices into one long α-helix that takes part in the interaction with MnmG. The physiological importance of this region, and of other regions within the MnmE-MnmG interface, is illustrated by the observation that mutations in corresponding residues of human GTPBP3-MTO1 are associated with disease.

**Figure 6:**
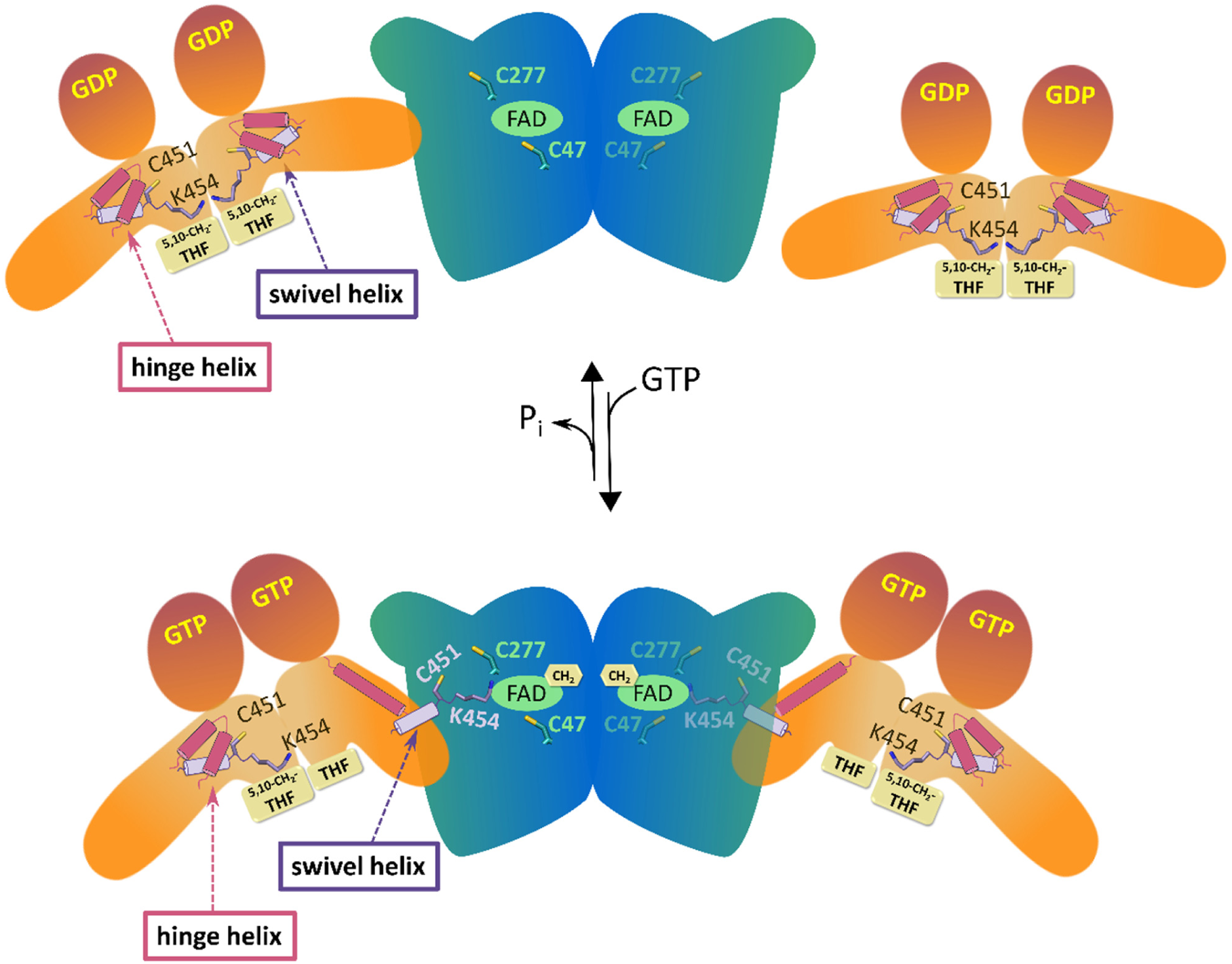
Schematic representation of the conformational changes that occur upon formation of the MnmE-MnmG complex. Large scale conformational changes in the interacting subunit(s) of MnmE are triggered by the structural rearrangement of the hinge helix. This translocates the swivel helix with its C-terminal K454 residue from the 5,10-CH_2_-THF binding pocket of MnmE to the FADH_2_ binding pocket of MnmE and assembles the three catalytic cysteine residues within the same pocket. We hypothesize that K454 could transport a methylene group from 5,10-CH_2_-THF to FADH_2_

The MnmE-MnmG complex catalyzes a highly sophisticated tRNA modification reaction orchestrated by GTP hydrolysis and involving three cofactors, with FAD and NADH binding to MnmG and 5,10-CH_2_-THF binding to MnmE [17]. Additionally, three cysteines are required for the reaction to occur, two provided by MnmG (C47 and C277) and one by MnmE (C451) [33]. One of the important remaining open questions was how all these cofactors, substrates and catalytic residues, provided by two different proteins, come together in an orchestrated fashion to add a carboxymethylaminomethyl (cmnm) group on the tRNA substrate [17]. In particular, the most recent findings propose a transfer of a methylene group from 5,10-CH_2_-THF to FADH_2_ resulting in a flavin iminium FADH[N^5^=CH2]^+^ intermediate as one of the first steps in the reaction mechanism [36,37]. In this mechanism FADH_2_ and 5,10-CH_2_-THF were proposed to form a covalent adduct during the transfer of the methylene group. However, in our structure of the α_2_β_2_ MnmEG complex, the distance between the N^5^ atom of FAD bound to the C subunit of MnmG and the N^5^ atom of a modelled THF cofactor in the B subunit of MnmE would be around 63 Å. This raises the question how the transfer of a methylene group would come about. The current cryo-EM structures now allow us to speculate on a plausible mechanism. In this scenario the MnmE C-terminal swivel helix and the highly conserved and functionally important ^450^FCIGK^454^ motif would play a crucial role. In the crystal structures of MnmE the swivel helix interacts with the MnmE N-terminal domain, and the ^450^FCIGK^454^ residues are inserted into the 5,10-CH_2_-THF binding pocket. In this orientation the side chain NH_2_-group of the C-terminal K454 residue is ideally positioned to perform a nucleophilic attack on the iminium tautomer of 5,10-CH_2_-THF leading to the transfer of the methylene group to K454 (**Fig. 5C**). Interaction with MnmG then induces a large structural rearrangement of the swivel helix, relocating it into the FADH_2_ binding pocket of MnmG (**Fig. 6**). In this orientation the methylene group could subsequently be transferred from K454 onto the N^5^ atom of FADH_2_ to form the FADH[N^5^=CH2]^+^ intermediate. Since in this conformation, the swivel helix covers the entrance to the MnmG FAD binding pocket, a final transfer of the methylene group to the substrate tRNA wobble uridine would next require another translocation of the swivel helix or of the entire MnmE molecule to allow the tRNA molecule to bind.

With respect to this proposed mechanism, it is also an intriguing observation that the interaction surface of MnmG for MnmE seems to overlap with the previously proposed tRNA binding surface. Indeed, the electrostatic potential surface of MnmG reveals that MnmE binds on a positively charged surface patch surrounding the FAD binding pocket of MnmG, and thereby also partially covers the entrance to the FAD pocket (**Fig. 4C**). Additionally, several residues of MnmG that are implicated in the interaction with MnmE, were previously found to affect tRNA binding [33]. This observation suggests that MnmE and tRNA might bind in a mutually exclusive way to MnmG. Alternatively, one could also imagine a more concerted mechanism, where one subunit of the MnmG dimer is bound to MnmE, while the other is bound to tRNA. In that respect it is interesting to note that only one tRNA molecule was found to bind the MnmG dimer in SAXS analysis [38]. In such a scenario the tRNA modification reaction would occur in a step-wise manner, where first the methylene group is transferred from 5,10-CH_2_-THF to FADH_2_ in the context of the MnmE-MnmG complex, after which the methylene group is transferred from FADH[N^5^=CH2]^+^ to tRNA U_34_ in the context of the MnmG-tRNA complex. Structures of either MnmG or the MnmE-MnmG complex bound to tRNA would be required to shed light on these questions.

## Data Availability

Cryo-EM density maps and structure coordinates have been deposited in the Electron Microscopy Data Bank (EMDB) and the Protein Data Bank (PDB), with accession codes EMD-52197, EMD-52198 and EMD-52199, and 9HIP, 9HIQ and 9HIR for the α_2_β_2_ complex, the full α_4_β_2_ complex and the MnmG-focused α_4_β_2_ complex, respectively. The simulation data are deposited in the Zenodo online repository under the identifier 10.5281/zenodo.14219035.

## Supplementary Data statement

Supplementary Data are available at NAR Online.

## Acknowledgements

W.V. would like to thank Fred Wittinghofer and Simon Meyer for sparking his interest in this topic and for the fantastic collaboration during the early stages of the work leading up to this publication. We would also like to thank Arjan Kortholt and Shehab Ismail for inspiring discussions throughout the entire project.

## Funding

This work was supported by a Strategic Research Program Financing from the VUB [SRP95]. D.B. was supported by the Research Foundation Flanders (FWO – grant G028821N). The computational resources and services used in this work were provided by the VSC (Flemish Supercomputer Center), funded by the Research Foundation Flanders (FWO) and the Flemish Government.

## Conflict of Interest Disclosure

The authors declare that they have no conflict of interest.

## Notes

https://doi.org/10.5281/zenodo.14219035

### Competing Interest Statement

The authors have declared no competing interest.

## References

1. El Yacoubi B, Bailly M, De Crécy-Lagard V (2012) Biosynthesis and function of posttranscriptional modifications of transfer RNAs. Annu Rev Genet 46: 69–95.

2. Motorin Y, Helm M (2010) tRNA stabilization by modified nucleotides. Biochemistry 49: 4934–4944.

3. Schimmel P (2018) The emerging complexity of the tRNA world: mammalian tRNAs beyond protein synthesis. Nat Rev Mol Cell Biol 19: 45–58.

4. Jackman JE, Alfonzo JD (2013) Transfer RNA modifications: nature’s combinatorial chemistry playground. Wiley Interdiscip Rev RNA 4: 35–48.

5. Gaik M, Kojic M, Wainwright BJ, Glatt S (2023) Elongator and the role of its subcomplexes in human diseases. EMBO Mol Med 15: e16418.

6. Torres AG, Batlle E, Ribas de Pouplana L (2014) Role of tRNA modifications in human diseases. Trends Mol Med 20: 306–314.

7. Santos M, Fidalgo A, Varanda AS, Oliveira C, Santos MAS (2019) tRNA Deregulation and Its Consequences in Cancer. Trends Mol Med 25: 853–865.

8. Arrondel C, Missoury S, Snoek R, Patat J, Menara G, Collinet B, Liger D, Durand D, Gribouval O, Boyer O, et al. (2019) Defects in t6A tRNA modification due to GON7 and YRDC mutations lead to Galloway-Mowat syndrome. Nat Commun 10: 3967.

9. Kirchner S, Ignatova Z (2015) Emerging roles of tRNA in adaptive translation, signalling dynamics and disease. Nat Rev Genet 16: 98–112.

10. Frye M, Harada BT, Behm M, He C (2018) RNA modifications modulate gene expression during development. Science 361: 1346–1349.

11. Endres L, Dedon PC, Begley TJ (2015) Codon-biased translation can be regulated by wobble-base tRNA modification systems during cellular stress responses. RNA Biol 12: 603–614.

12. Thongdee N, Jaroensuk J, Atichartpongkul S, Chittrakanwong J, Chooyoung K, Srimahaeak T, Chaiyen P, Vattanaviboon P, Mongkolsuk S, Fuangthong M (2019) TrmB, a tRNA m7G46 methyltransferase, plays a role in hydrogen peroxide resistance and positively modulates the translation of katA and katB mRNAs in Pseudomonas aeruginosa. Nucleic Acids Res 47: 9271– 9281.

13. Agris PF, Eruysal ER, Narendran A, Väre VYP, Vangaveti S, Ranganathan S V. (2018) Celebrating wobble decoding: Half a century and still much is new. RNA Biol 15: 537–553.

14. Suzuki T (2021) The expanding world of tRNA modifications and their disease relevance. Nat Rev Mol Cell Biol 22: 375–392.

15. Yokoyama S, Watanabe T, Murao K, Ishikura H, Yamaizumi Z, Nishimura S, Miyazawa T (1985) Molecular mechanism of codon recognition by tRNA species with modified uridine in the first position of the anticodon. Proc Natl Acad Sci U S A 82: 4905–4909.

16. Armengod ME, Moukadiri I, Prado S, Ruiz-Partida R, Benítez-Páez A, Villarroya M, Lomas R, Garzón MJ, Martínez-Zamora A, Meseguer S, et al. (2012) Enzymology of tRNA modification in the bacterial MnmEG pathway. Biochimie 94: 1510–1520.

17. Fislage M, Wauters L, Versées W (2016) Invited review: MnmE, a GTPase that drives a complex tRNA modification reaction. Biopolymers 105: 568–579.

18. Moukadiri I, Prado S, Piera J, Velázquez-campoy A, Björk GR, Armengod ME (2009) Evolutionarily conserved proteins MnmE and GidA catalyze the formation of two methyluridine derivatives at tRNA wobble positions. Nucleic Acids Res 37: 7177–7193.

19. Suzuki T, Suzuki T, Wada T, Saigo K, Watanabe K (2002) Taurine as a constituent of mitochondrial tRNAs: new insights into the functions of taurine and human mitochondrial diseases. EMBO J 21: 6581–6589.

20. Jaciuk M, Scherf D, Kaszuba K, Gaik M, Rau A, Kościelniak A, Krutyhołowa R, Rawski M, Indyka P, Graziadei A, et al. (2023) Cryo-EM structure of the fully assembled Elongator complex. Nucleic Acids Res 51: 2011–2032.

21. Charif M, Titah SMC, Roubertie A, Desquiret-Dumas V, Gueguen N, Meunier I, Leid J, Massal F, Zanlonghi X, Mercier J, et al. (2015) Optic neuropathy, cardiomyopathy, cognitive disability in patients with a homozygous mutation in the nuclear MTO1 and a mitochondrial MT-TF variant. Am J Med Genet A 167A: 2366–2374.

22. Kopajtich R, Nicholls TJ, Rorbach J, Metodiev MD, Freisinger P, Mandel H, Vanlander A, Ghezzi D, Carrozzo R, Taylor RW, et al. (2014) Mutations in GTPBP3 cause a mitochondrial translation defect associated with hypertrophic cardiomyopathy, lactic acidosis, and encephalopathy. Am J Hum Genet 95: 708–720.

23. Ghezzi D, Baruffini E, Haack TB, Invernizzi F, Melchionda L, Dallabona C, Strom TM, Parini R, Burlina AB, Meitinger T, et al. (2012) Mutations of the mitochondrial-tRNA modifier MTO1 cause hypertrophic cardiomyopathy and lactic acidosis. Am J Hum Genet 90: 1079–1087.

24. Shippy DC, Eakley NM, Bochsler PN, Fadl A a. (2011) Biological and virulence characteristics of Salmonella enterica serovar Typhimurium following deletion of glucose-inhibited division (gidA) gene. Microb Pathog 50: 303–313.

25. Sha J, Kozlova E V., Fadl AA, Olano JP, Houston CW, Peterson JW, Chopra AK (2004) Molecular characterization of a glucose-inhibited division gene, gidA, that regulates cytotoxic enterotoxin of Aeromonas hydrophila. Infect Immun 72: 1084–1095.

26. Srimahaeak T, Thongdee N, Chittrakanwong J, Atichartpongkul S, Jaroensuk J, Phatinuwat K, Phaonakrop N, Jaresitthikunchai J, Roytrakul S, Mongkolsuk S, et al. (2023) Pseudomonas aeruginosa GidA modulates the expression of catalases at the posttranscriptional level and plays a role in virulence. Front Microbiol 13: 1079710.

27. Scrima A, Vetter IR, Armengod ME, Wittinghofer A (2005) The structure of the TrmE GTP-binding protein and its implications for tRNA modification. EMBO J 24: 23–33.

28. Meyer S, Böhme S, Krüger A, Steinhoff HJ, Klare JP, Wittinghofer A (2009) Kissing G domains of MnmE monitored by X-ray crystallography and pulse electron paramagnetic resonance spectroscopy. PLoS Biol 7: e1000212.

29. Gasper R, Meyer S, Gotthardt K, Sirajuddin M, Wittinghofer A (2009) It takes two to tango: regulation of G proteins by dimerization. Nat Rev Mol Cell Biol 10: 423–429.

30. Scrima A, Wittinghofer A (2006) Dimerisation-dependent GTPase reaction of MnmE: how potassium acts as GTPase-activating element. EMBO J 25: 2940–2951.

31. Meyer S, Wittinghofer A, Versées W (2009) G-Domain Dimerization Orchestrates the tRNA Wobble Modification Reaction in the MnmE/GidA Complex. J Mol Biol 392: 910–922.

32. Meyer S, Scrima A, Versées W, Wittinghofer A (2008) Crystal Structures of the Conserved tRNA-Modifying Enzyme GidA: Implications for Its Interaction with MnmE and Substrate. J Mol Biol 380: 532–547.

33. Osawa T, Ito K, Inanaga H, Nureki O, Tomita K, Numata T (2009) Conserved Cysteine Residues of GidA Are Essential for Biogenesis of 5-Carboxymethylaminomethyluridine at tRNA Anticodon. Structure 17: 713–724.

34. Shi R, Villarroya M, Ruiz-Partida R, Li Y, Proteau A, Prado S, Moukadiri I, Benítez-Páez A, Lomas R, Wagner J, et al. (2009) Structure-function analysis of Escherichia coli MnmG (GidA), a highly conserved tRNA-modifying enzyme. J Bacteriol 191: 7614–7619.

35. Yim L, Moukadiri I, Björk GR, Armengod ME (2006) Further insights into the tRNA modification process controlled by proteins MnmE and GidA of Escherichia coli. Nucleic Acids Res 34: 5892–5905.

36. Bommisetti P, Young A, Bandarian V (2022) Elucidation of the substrate of tRNA-modifying enzymes MnmEG leads to in vitro reconstitution of an evolutionarily conserved uridine hypermodification. J Biol Chem 298: 102548.

37. Bommisetti P, Bandarian V (2023) Insights into the Mechanism of Installation of 5-Carboxymethylaminomethyl Uridine Hypermodification by tRNA-Modifying Enzymes MnmE and MnmG. J Am Chem Soc 145: 26947–26961.

38. Fislage M, Brosens E, Deyaert E, Spilotros A, Pardon E, Loris R, Steyaert J, Garcia-Pino A, Versées W (2014) SAXS analysis of the tRNA-modifying enzyme complex MnmE/MnmG reveals a novel interaction mode and GTP-induced oligomerization. Nucleic Acids Res 42: 5978–5992.

39. Bepler T, Morin A, Rapp M, Brasch J, Shapiro L, Noble AJ, Berger B (2019) Positive-unlabeled convolutional neural networks for particle picking in cryo-electron micrographs. Nat Methods 16: 1153–1160.

40. Terwilliger TC, Ludtke SJ, Read RJ, Adams PD, Afonine P V. (2020) Improvement of cryo-EM maps by density modification. Nat Methods 17: 923–927.

41. Adams PD, Afonine P V., Bunkóczi G, Chen VB, Davis IW, Echols N, Headd JJ, Hung LW, Kapral GJ, Grosse-Kunstleve RW, et al. (2010) PHENIX: a comprehensive Python-based system for macromolecular structure solution. Acta Crystallogr D Biol Crystallogr 66: 213–221.

42. Jumper J, Evans R, Pritzel A, Green T, Figurnov M, Ronneberger O, Tunyasuvunakool K, Bates R, Žídek A, Potapenko A, et al. (2021) Highly accurate protein structure prediction with AlphaFold. Nature 596: 583–589.

43. Pettersen EF, Goddard TD, Huang CC, Meng EC, Couch GS, Croll TI, Morris JH, Ferrin TE (2021) UCSF ChimeraX: Structure visualization for researchers, educators, and developers. Protein Sci 30: 70–82.

44. Emsley P, Lohkamp B, Scott WG, Cowtan K (2010) Features and development of Coot. Acta Crystallogr Sect D Biol Crystallogr 66: 486–501.

45. Huang J, Rauscher S, Nawrocki G, Ran T, Feig M, De Groot BL, Grubmüller H, MacKerell AD (2017) CHARMM36m: an improved force field for folded and intrinsically disordered proteins. Nat Methods 14: 71–73.

46. Abraham MJ, Murtola T, Schulz R, Páll S, Smith JC, Hess B, Lindah E (2015) GROMACS: High performance molecular simulations through multi-level parallelism from laptops to supercomputers. SoftwareX 1–2: 19–25.

47. Bussi G, Donadio D, Parrinello M (2007) Canonical sampling through velocity rescaling. J Chem Phys 126: 014101.

48. Svergun D, Barberato C, Koch MH (1995) CRYSOL - A program to evaluate X-ray solution scattering of biological macromolecules from atomic coordinates. J Appl Crystallogr 28: 768– 773.

49. Petoukhov M V., Franke D, Shkumatov A V., Tria G, Kikhney AG, Gajda M, Gorba C, Mertens HDT, Konarev P V., Svergun DI (2012) New developments in the ATSAS program package for small-angle scattering data analysis. J Appl Crystallogr 45: 342–350.

50. Böhme S, Meyer S, Krüger A, Steinhoff HJ, Wittinghofer A, Klare JP (2010) Stabilization of G domain conformations in the tRNA-modifying MnmE-GidA complex observed with double electron electron resonance spectroscopy. J Biol Chem 285: 16991–17000.

51. Prado S, Villarroya M, Medina M, Armengod ME (2013) The tRNA-modifying function of MnmE is controlled by post-hydrolysis steps of its GTPase cycle. Nucleic Acids Res 41: 6190–6208.

52. Krissinel E, Henrick K (2007) Inference of Macromolecular Assemblies from Crystalline State. J Mol Biol 372: 774–797.

53. Kamps R, Szklarczyk R, Theunissen TE, Hellebrekers DMEI, Sallevelt SCEH, Boesten IB, De Koning B, Van Den Bosch BJ, Salomons GS, Simas-Mendes M, et al. (2018) Genetic defects in mtDNA-encoded protein translation cause pediatric, mitochondrial cardiomyopathy with early-onset brain disease. Eur J Hum Genet 26: 537–551.

54. Monda E, Lioncino M, Caiazza M, Simonelli V, Nesti C, Rubino M, Perna A, Mauriello A, Budillon A, Pota V, et al. (2023) Clinical, Genetic, and Histological Characterization of Patients with Rare Neuromuscular and Mitochondrial Diseases Presenting with Different Cardiomyopathy Phenotypes. Int J Mol Sci 24: 9108.

55. O’Byrne JJ, Tarailo-Graovac M, Ghani A, Champion M, Deshpande C, Dursun A, Ozgul RK, Freisinger P, Garber I, Haack TB, et al. (2018) The genotypic and phenotypic spectrum of MTO1 deficiency. Mol Genet Metab 123: 28–42.

56. Zhou C, Wang J, Zhang Q, Yang Q, Yi S, Shen Y, Luo J, Qin Z (2022) Clinical and genetic analysis of combined oxidative phosphorylation defificiency-10 caused by MTO1 mutation. Clin Chim Acta 526: 74–80.

