## Supplementary Material for "Cryo-EM structures of the MnmE-MnmG complex reveal large conformational changes and provide new insights into the mechanism of tRNA modification"

#### SUPPLEMENTARY FIGURES

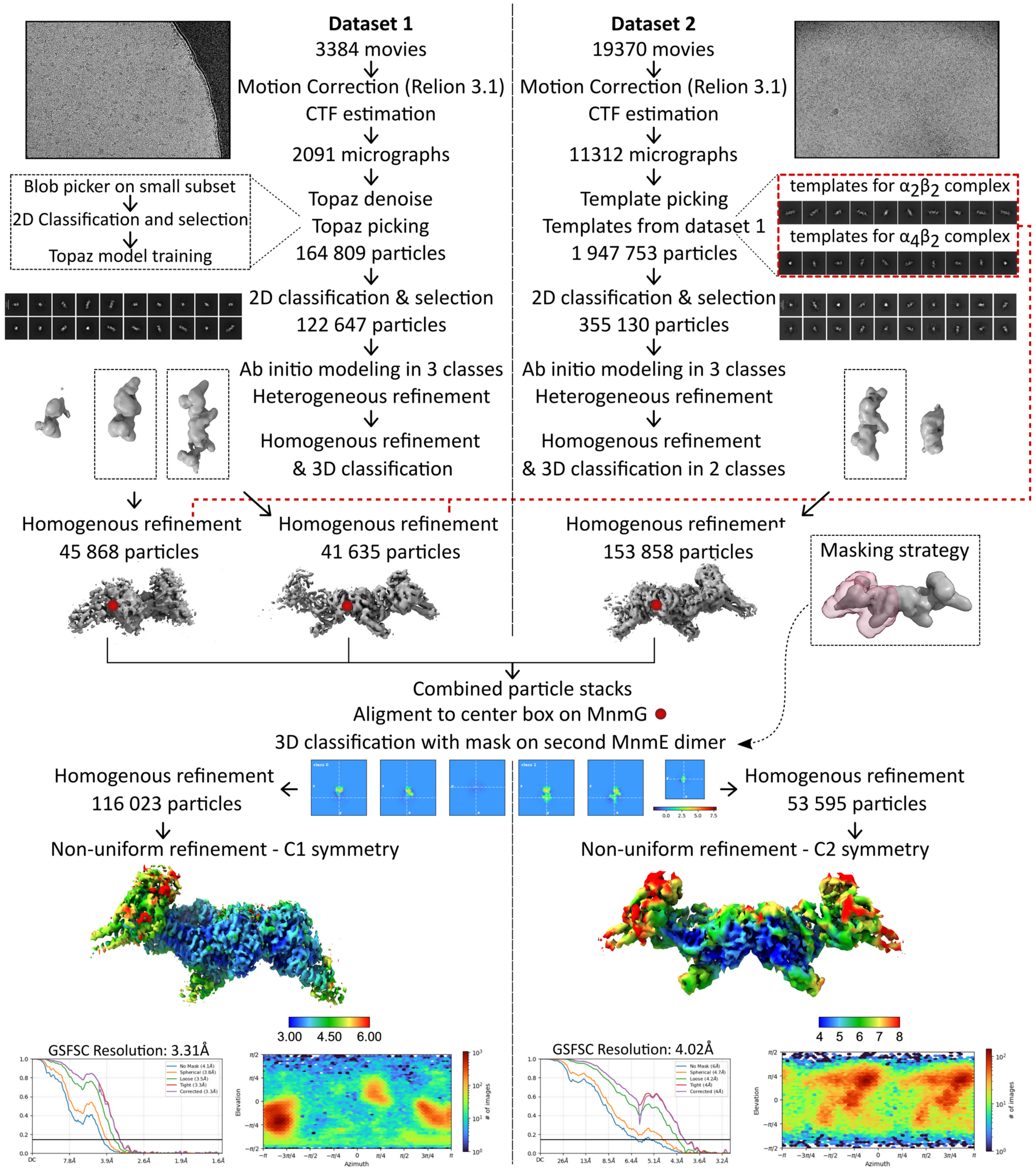

**Figure S1. Cryo-electron microscopy (cryo-EM) workflow to obtain the maps corresponding to the $\alpha_2\beta_2$ and $\alpha_4\beta_2$ MnmEG complexes.** Flow chart of the cryo-EM data processing for both datasets, including picking strategies, 2D classification, 3D classification with mask and density map reconstructions. Local resolution maps, Fourier shell correlation (FSC) curves, particle distribution plots, and cryo-EM density maps are also shown. All analyses were performed using cryoSPARC v3.3, unless stated otherwise. Details are provided in the 'Materials and methods' section.

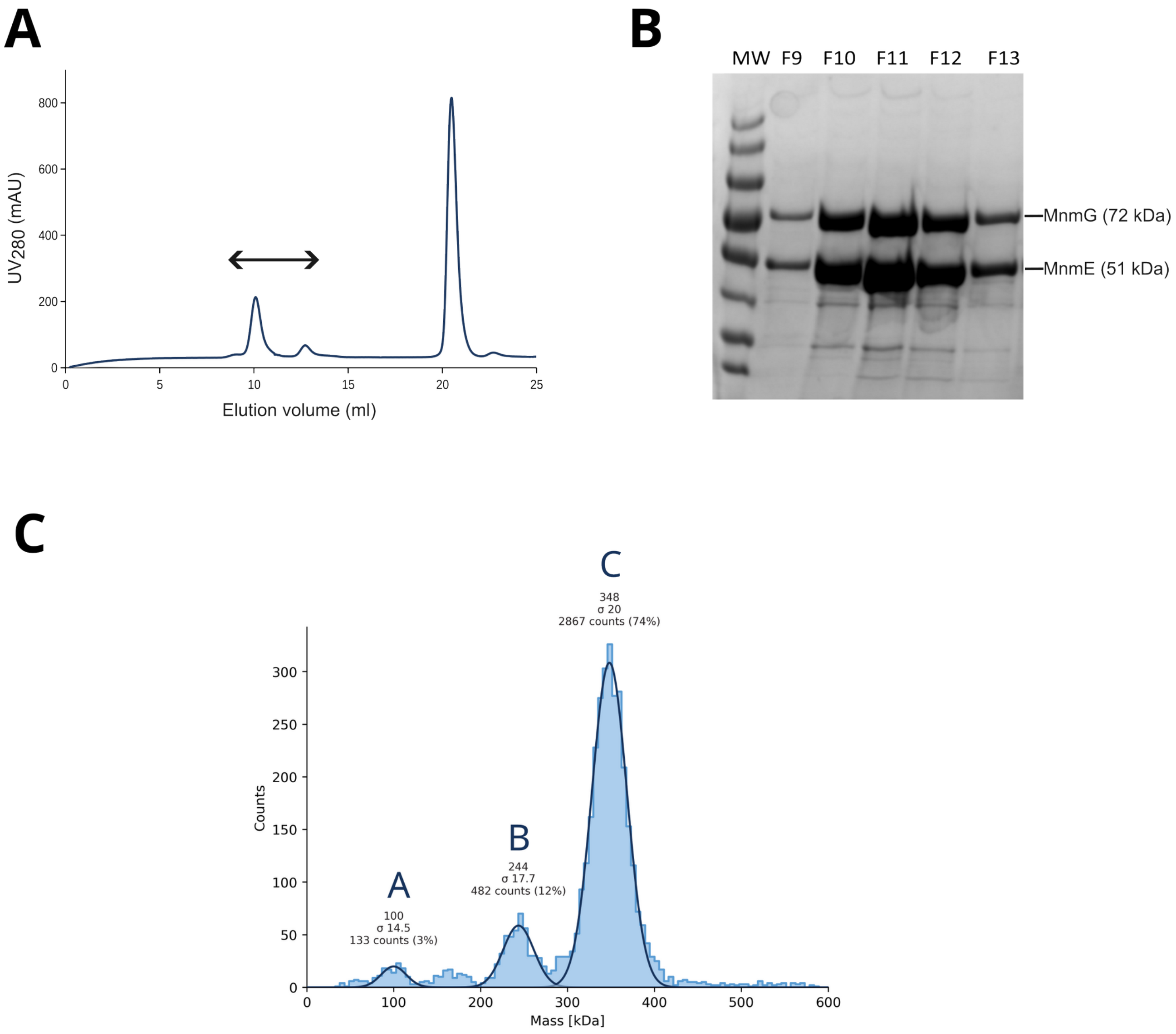

| Peak | Molecular weight (kDa) | Molar fraction | Counts | Distribution (σ) |
|---|---|---|---|---|
| A | 100 | 0.03 | 133 | 14.5 |
| B | 244 | 0.12 | 482 | 17.7 |
| C | 348 | 0.74 | 2867 | 20 |

**Figure S2. Purification and analysis of the MnmEG complex. (A)** Purification of the MnmEG complex on size exclusion chromatography (SEC). 100 µM MnmE and 50 µM MnmG were mixed in the presence of 1 mM GppNHp and 1 mM FAD, and the mixture was loaded on a Superdex 200 10/300 column to separate the $\alpha_4\beta_2$ complex. Fractions that were analyzed on SDS-PAGE are indicated with an arrow. **(B)** SDS-PAGE analysis of the peak fractions from SEC corresponding to the $\alpha_4\beta_2$ complex. Fraction 10 and 11 (F10-F11) were pooled for analysis by mass photometry. (**C**) Mass photometry analysis of the pooled fractions 10-11 at a concentration of 100 nM on a Refeyn OneMP instrument. This analysis indicates that more than 70% of the population attains a molecular mass close to the 348 kDa expected for the $\alpha_4\beta_2$ complex.

**A**

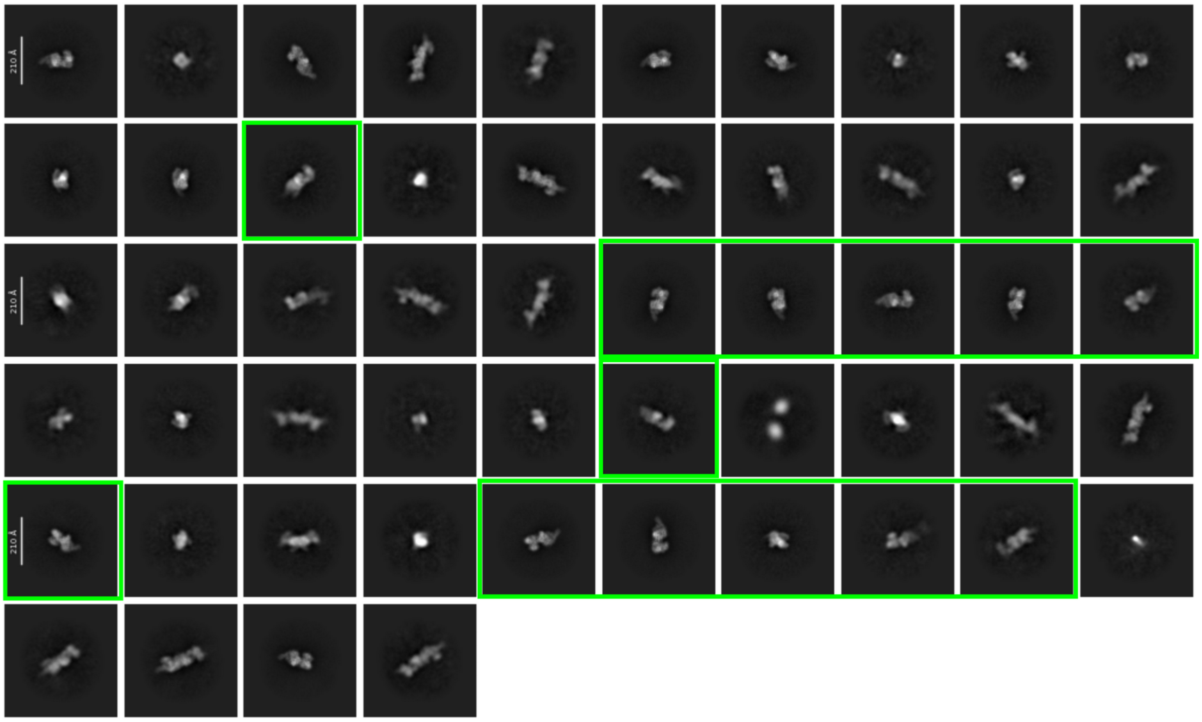

**B**

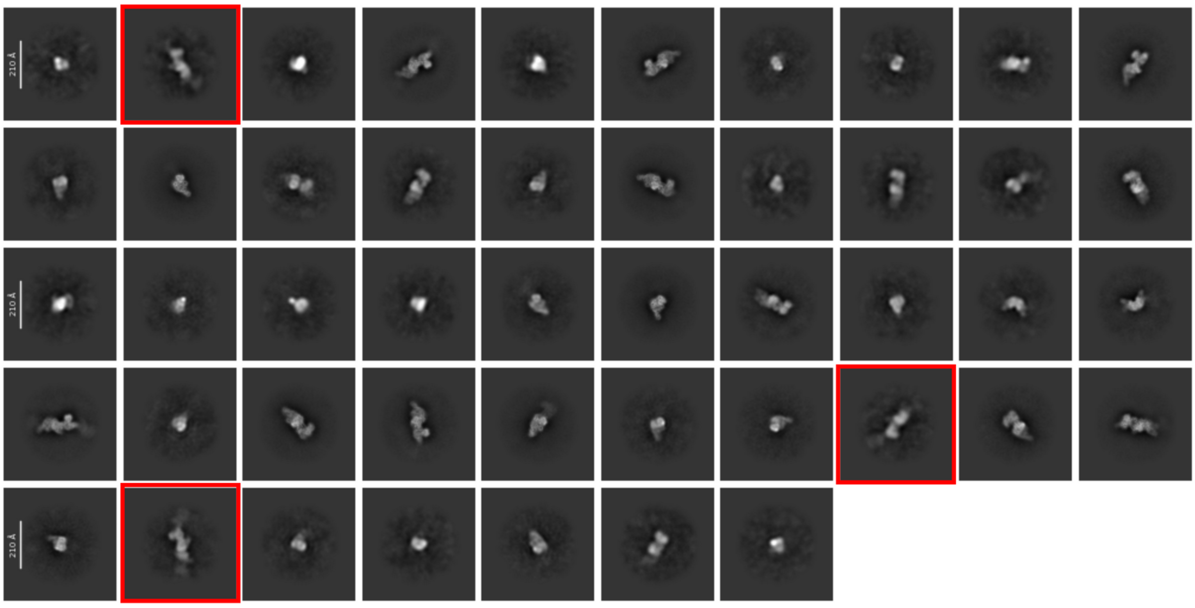

**Figure S3. Representative 2D classes of the MnmEG complexes in the two cryo-EM datasets that were collected. (A)** 2D classes from particles extracted in dataset 1 with a box size of 504 Å. Most 2D classes represent very elongated particles corresponding to the $\alpha_4\beta_2$ complex. Nevertheless, in some classes (indicated with a green square) a slightly less elongated particle is observed, indicating that also the smaller $\alpha_2\beta_2$ oligomeric state is present. **(B)** 2D classes from particles extracted in dataset 2 with a box size of 504 Å. Although most classes represent the smaller $\alpha_2\beta_2$ oligomeric state, some classes display a blurred density representing the presence of a second MnmE dimer in the $\alpha_4\beta_2$ complex (indicated by red squares)

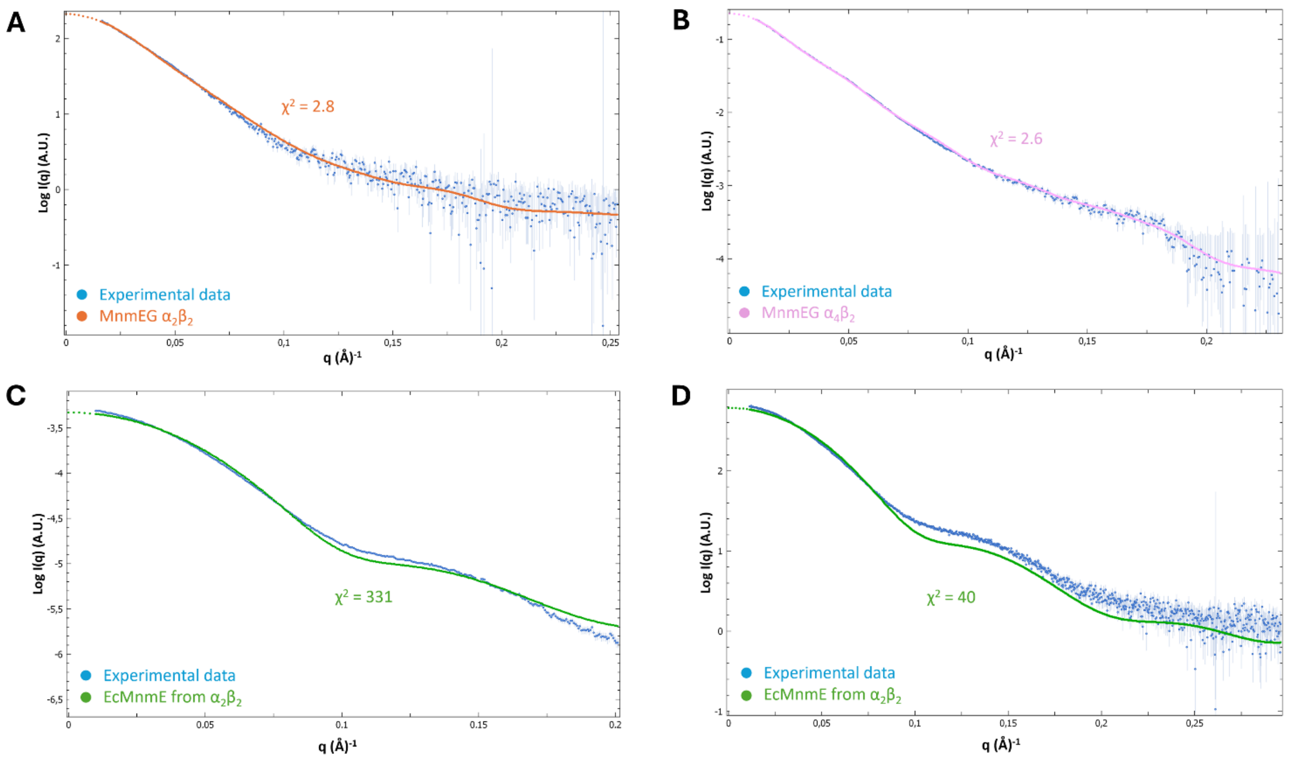

**Figure S4. Comparison of the cryo-EM models of the $\alpha_2\beta_2$ and $\alpha_4\beta_2$ complexes with prior obtained solution SAXS data using Crysol** (*Fislage et al. Nucleic Acids Res. 42(9):5978-92, 2014*). **(A)** Superposition of experimentally obtained SAXS data for the $\alpha_2\beta_2$ complex (blue) and the theoretical scatter curve recalculated from the cryo-EM model of the $\alpha_2\beta_2$ complex (red). The experimental scattering curve agrees well with the theoretical scattering curves obtained from the model, with a $\chi 2$ value of 2.8. **(B)** Superposition of experimentally obtained SAXS data for the $\alpha_4\beta_2$ complex (blue) and the theoretical scatter curve recalculated from the cryo-EM model of the $\alpha_4\beta_2$ complex (purple). The experimental scattering curve agrees well with the theoretical scattering curves obtained from the model, with a $\chi 2$ value of 2.6. **(C/D)** Superposition of the theoretical scatter curve calculated from the asymmetric MnmE dimer extracted from the $\alpha_2\beta_2$ complex (green) and the experimentally obtained SAXS data (blue) for MnmE bound to GDP*AlF (**C**) or GppNHp (**D**). This analysis shows significant differences in the overall shape of GppNHp- or GDP*AlF4-bound MnmE in solution and the conformation of MnmE in complex with MnmG. For all details on the conditions of the experimentally obtained scatter curves see: *Fislage et al. Nucleic Acids Res. 42(9):5978-92, 2014*.

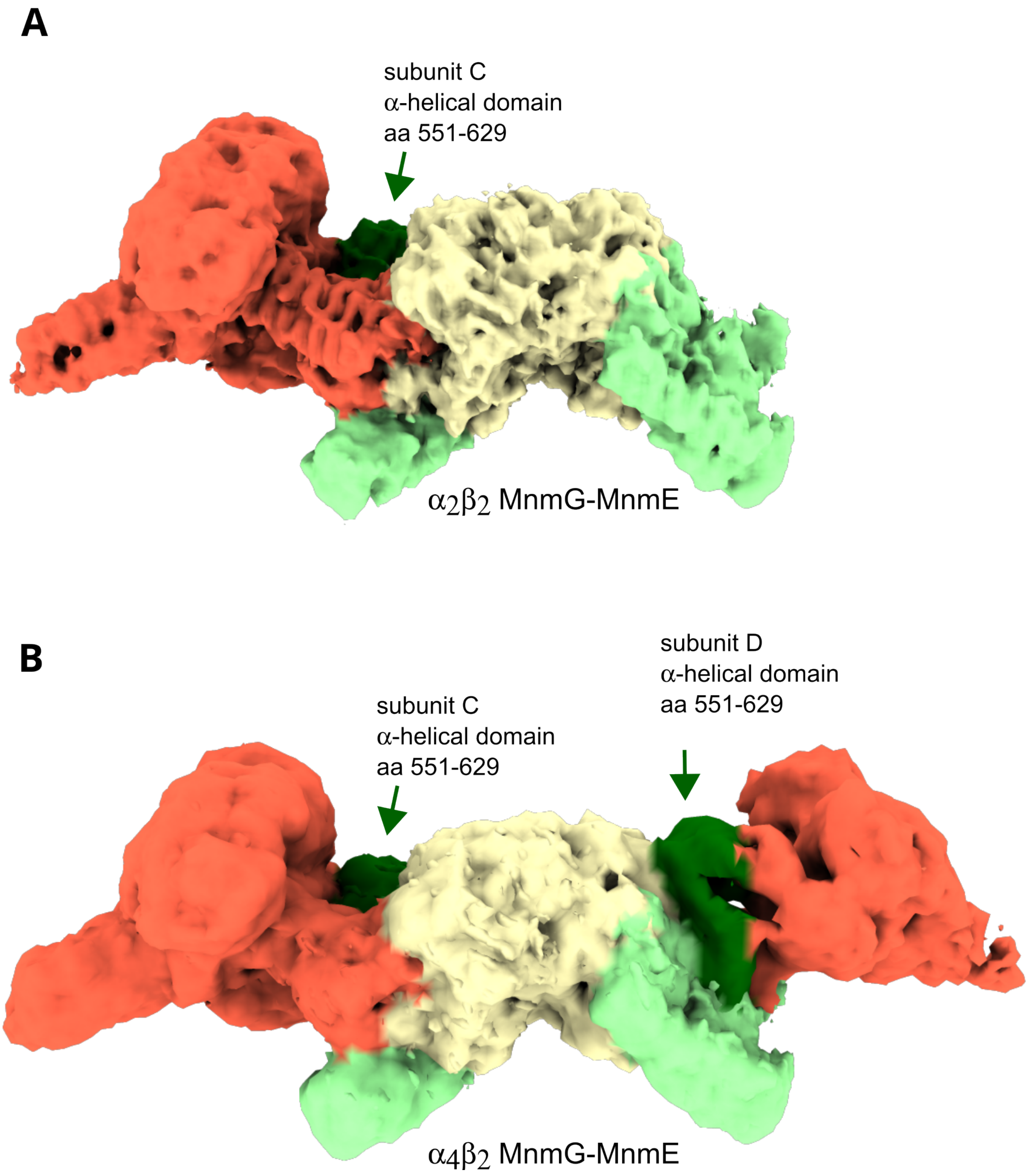

**Figure S5. Cryo-EM maps of the MnmEG $\alpha_2\beta_2$ and $\alpha_4\beta_2$ complexes without imposing symmetry**. The maps of MnmE are colored red. The map of MnmG is colored yellow, with the N-terminal part of its α-helical domains colored pale green and the C-terminal part of its a-helical domains (551-629) colored dark green. Both maps represent full reconstructions without any imposed symmetry.

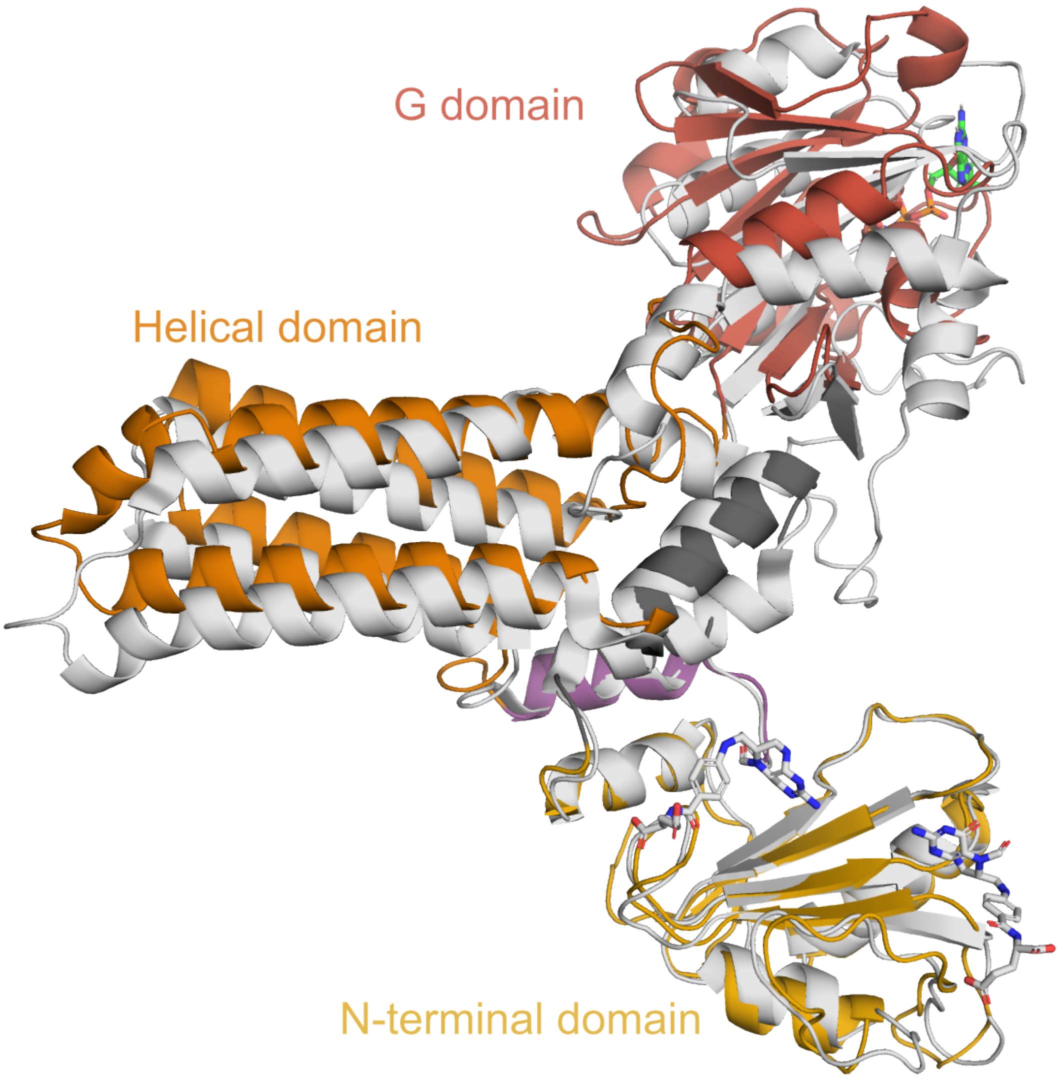

**Figure S6. Superposition of the A subunit of MnmE extracted from the $\alpha_2\beta_2$ complex on a subunit of the crystal structure of MnmE from *Thermatoga maritima*** (TmMnmE, PDB 1XZQ). The domains of the A subunit of MnmE from the $\alpha_2\beta_2$ complex are indicated with the N-terminal domain, helical domain and G domain in yellow, orange and red, respectively, while the hinge helix and swivel helix are shown in dark grey and purple, respectively. The subunit of TmMnmE is shown in grey.

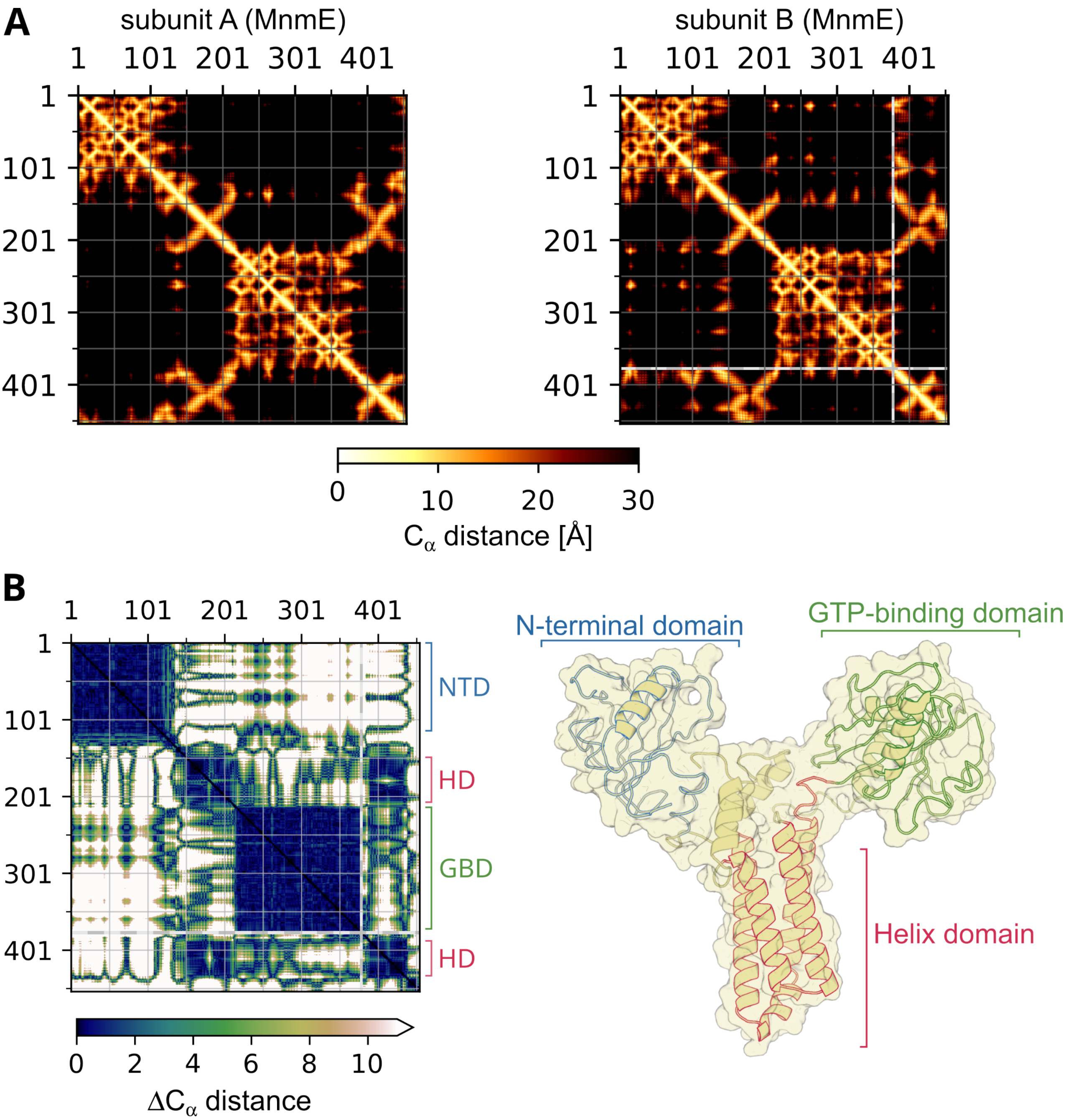

**Figure S7**: **Analysis and comparison of the Cα distance matrices of the MnmE subunits A and B in the $\alpha_2\beta_2$ model.** (**A**) Cα distance matrix of MnmE subunit A (*left*) and subunit B (*right*). Unresolved residues are shown in a grey checkerboard pattern. (**B**) Difference in the Cα-Cα distance matrices between the subunit A and subunit B conformations of MnmE (*left*) and mapping of the domains that act as rigid bodies on the subunit structure of MnmE. It can be observed that the N-terminal domain (NTD), G domain (GBD) and the four-helix bundle of the helical domain (HD) move relative to each other but retain their internal conformation.

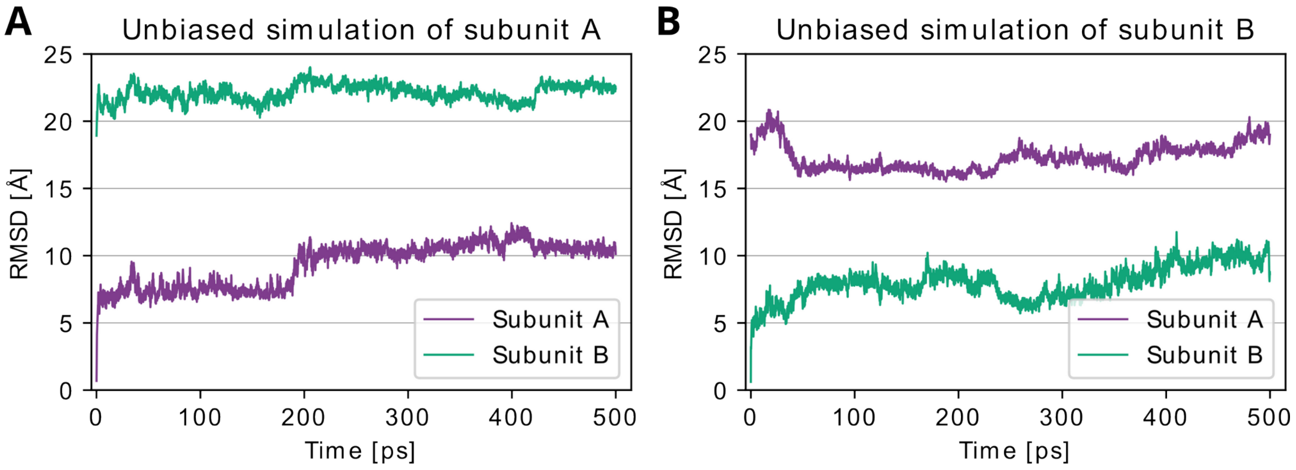

**Figure S8. RMSD values of the unbiased MD simulations of monomeric MnmE in relation to subunit A (violet) and subunit B (teal).** (**A**) Simulation that was started from the subunit A conformation. (**B**) Simulation that was started from the subunit B conformation.

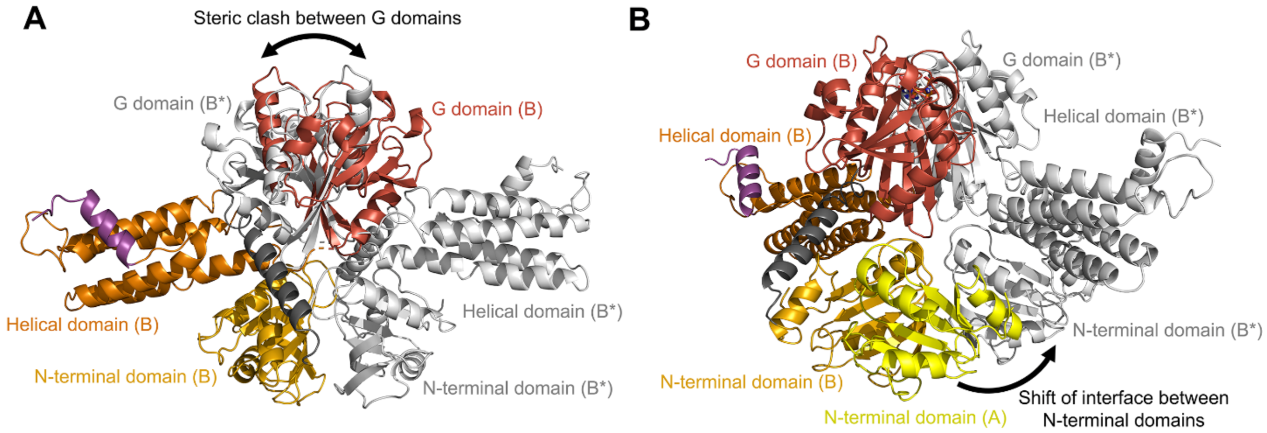

**Figure S9. MnmE is unlikely to form a symmetrical dimer, in which both subunits adopt a conformation disposed to interact with MnmG ("B subunit conformation"), within the $\alpha_2\beta_2$ or $\alpha_4\beta_2$ complexes.** (**A**) Creation of a symmetrical B-B* MnmE dimer, while maintaining the interface between the N-terminal domains, would result in severe clashes of the G-domains of the adjacent subunits. (**B**) Creation of a symmetrical B-B* MnmE dimer, while maintaining the interface between the G domains, would require a completely different interaction interface between the N-terminal domains of the constitutive MnmE dimer. The domains of the MnmE subunit B are colored similarly to Figure 3A. The symmetry variant of subunit B (B*) is shown in grey. In panel (B), the N-terminal domain of MnmE subunit A is shown in yellow.

### SUPPLEMENTARY TABLES

**Table S1.** Cryo-EM data collection, refinement and validation statistics

| | MnmE-MnmG $\alpha_2\beta_2$ (EMD-52197) (PDB 9HIP) | MnmE-MnmG $\alpha_4\beta_2$ (EMD-52198) (PDB 9HIQ) | MnmG focused $\alpha_4\beta_2$ (EMD-52199) (PDB 9HIR) |
|---|---|---|---|
| **Data collection** | | | |
| Microscope | CryoARM300 | | |
| Voltage (kV) | 300 | | |
| Electron exposure (e–/Å$^2$) | 62 | | |
| Energy filter slit width | 20 eV | | |
| Detector | Gatan K3 | | |
| Magnification | 60,000 | | |
| Defocus range (μm) | 0.5 – 3 | | |
| Pixel size (Å) | 0.7596 | | |
| Initial particles (no.) | 2,112,562 | | |
| Symmetry imposed | C1 | C2 | C1 |
| Final particles (no.) | 116,023 | 53,595 | 107,190 |
| Map mean resolution (Å) | 3.31 | 4.02 | 4.12 |
| FSC threshold | 0.143 | 0.143 | 0.143 |
| **Refinement** | | | |
| Model composition | | | |
| Non-hydrogen atoms | 15820 | 23248 | 9774 |
| Protein residues | 2043 | 3029 | 1244 |
| Ligands | 3 | 6 | 2 |
| R.m.s. deviations | | | |
| Bond lengths (Å) | 0.002 | 0.004 | 0.004 |
| Bond angles (°) | 0.416 | 0.532 | 0.673 |
| Validation | | | |
| MolProbity score | 2.53 | 2.33 | 1.9 |
| Clash score | 43.87 | 34.4 | 13.49 |
| Poor rotamers (%) | 0.55 | 0 | 0 |
| Ramachandran plot | | | |
| Favored (%) | 93.94 | 95.57 | 96.12 |
| Allowed (%) | 5.96 | 4.43 | 3.88 |
| Disallowed (%) | 0.10 | 0 | 0 |
| *B* factors (Å$^2$) | | | |
| Protein | 63.15 | 160.04 | 70.92 |
| Ligand | 70.73 | 165.52 | 65.03 |
